# YnaI exemplifies the diversity of structural gating mechanisms in mechanosensitive channels of small conductance

**DOI:** 10.1101/2024.12.30.628871

**Authors:** Vanessa J. Flegler, Akiko Rasmussen, Rainer Hedrich, Tim Rasmussen, Bettina Böttcher

## Abstract

Osmotically varying environments are challenging for bacterial cells. Sudden drops in osmolytes cause an increased membrane tension and rupture the cells in the absence of protective mechanisms. One family of protective proteins are mechanosensitive channels of small conductance that open in response to membrane tension. Although these channels have a common architecture, they vary widely in the number of transmembrane helices, conductivity, and gating characteristics. Despite of several structures of channels in the open and closed state, the underlying common principles of the gating mechanism are not well understood. Here we show that YnaI opens by radial relocation of the transmembrane sensor paddles together with a shortening of the pore. This contrasts the prototypic smaller MscS which tilts the sensor paddles and retains the pore length. A chimera of both channels with the YnaI sensor paddles and the pore containing C-terminal part of MscS has the conductivity of the pore donor and the tension response of the paddle donor together with the conformational changes of the respective donor. Our research shows that elements with different types of structural rearrangements can be mixed and matched within one channel as long as they support the common area expansion on the periplasmic side.

## Introduction

Bacteria are exposed to many changes in their environment and must adapt quickly to survive. One of these challenges is the maintenance of osmotic homeostasis. A hypoosmotic shock will rapidly drive water into the cell, eventually tensing the membrane up to the point of rupture. To prevent cell lysis, bacteria have mechanosensitive channels that are gated in response to the elevated membrane tension and release solutes restoring the turgor within the cells (Martinac *et al*, 1987; Levina *et al*, 1999; Booth & Blount, 2012). Different types of mechanosensitive channels convey a graded response mechanism. One family is designated as mechanosensitive channels of small conductance because of their electrophysiological properties. Typically, they open at membrane tensions much lower than the point of rupture but release only small solutes with a very small flux. One of the first discovered family members is MscS (Martinac *et al*, 1987), which is a homoheptamer comprised of a cytosolic vestibule and a transmembrane part with a central pore surrounded by seven sensor paddles. In addition, *Escherichia coli* (*E. coli*) has five larger sized paralogues with a similar overall architecture but more transmembrane (TM) helices and different gating characteristics (Edwards *et al*, 2012; Schumann *et al*, 2010; Li *et al*, 2002).

One of the medium sized paralogues is YnaI, with 5 TM helices. YnaI opens at a larger membrane tension and with a smaller conductivity than all other MscS-like paralogues in *E. coli*. Therefore, its importance for the protection against osmotic challenges is enigmatic as only few solutes will leave the cell via YnaI in the presence of the other paralogues. Structural studies have shown that the main difference between YnaI and MscS are extended sensor paddles, which create a periplasmic indentation in YnaI, and much smaller portals in the vestibule that provide the cytoplasmic access to the pore. These lateral portals convey selectivity and restrict the conductance of some channels In YnaI the portals are particularly small and limit the maximal conductance to only 0.1 nS compared to 1.3 nS in MscS. The arrangement of vestibule, pore and paddles creates hydrophobic pockets which extend into the cytosolic space, where they are populated by lipid molecules in an orientation almost parallel to the plane of membrane (Flegler *et al*, 2020; Hu *et al*, 2021; Catalano *et al*, 2021). The TM paddles are in contact with the surrounding membrane and are therefore responsible for sensing the membrane tension, while the vestibule with the pore formed by the TM3 helices convey conductivity and selectivity.

Opening of MscS-like channels is characterised by an outwards movement of the sensor paddles on the periplasmatic side, resulting in an increased cross section in the membrane plane (Wang *et al*, 2008; Deng *et al*, 2020; Mount *et al*, 2022). The paddle rearrangement pulls the pore helices apart and enables the passage of small solutes and water molecules between the vestibule and the periplasm. In the prototypic MscS the paddle movement is concomitant with tilting of the whole paddle and relocation of the pore into the plane of membrane. The movement of the paddles is accompanied by a reorientation of the lipids in the pocket and a transition from a curved local membrane environment into a planar bilayer arrangement in the open state. A hallmark of the tilting mechanism is a very long outer paddle helix, which extends into the cytosol and forms the entrance to the cytosolic pockets. In the closed state these helices come together on the periplasmic side, resulting in a cone shaped TM part, and trap the head group of a so-called hook or gate keeper lipid in the centre of the membrane (Reddy *et al*, 2019; Rasmussen *et al*, 2019). Alignment of lipid molecules in the membrane with the hook lipid induces the local curvature in the surrounding membrane (Park *et al*, 2023).

These observations have raised the expectancy that all mechanosensitive channels open with a similar mechanism. However, YnaI is lacking some of the structural key elements: In the closed state, the channel is less cone-shaped, and the paddles interdigitate with the lipids in the membrane without inducing marked local curvature. Furthermore, the outermost paddle helix is short and buried in the membrane, which makes tilting unlikely without distorting the local membrane environment. With electron cryo-microscopy and image processing we show that the underlying mechanism for channel opening in YnaI is distinct from that of MscS. A chimera of both channels shows that the sensor paddles dictate the structural rearrangements while the pore/vestibule module dictates the conductivity.

## Results

### Lipid supplementation stabilises YnaI in the closed state

Closed YnaI was purified for structure determination by electron cryo-microscopy in n-dodecyl-β-maltoside (DDM) supplemented with 0.5 mg/ml Azolectin (condition I+), resulting in a 2.2 Å map (figure S1, figure S2, figure S3). The model built from residues 3-334 resembled the one of YnaI purified in LMNG by (Hu *et al*, 2021) with minor differences in the outermost TM helix (TM(-2)), which is frame shifted by 3 residues In our model Trp29 in TM(-2) is at the cytosolic interface of the membrane. This is the only tryptophan in the sensor paddle of YnaI and provides a typical side chain found in the amphiphilic region of membranes where it tightly anchors the TM helix (De Jesus & Allen, 2013). Each sensor paddle has four TM helices that are interconnected by 1-2 hydrogen bonds to the adjacent paddle helix. For example, Asn8 in TM(-2) makes a hydrogen bond with Tyr63 in TM(-1). This hydrogen bonding network suggests that the whole paddle forms a rigid entity within the membrane (figure 1a).

**Figure 1:**
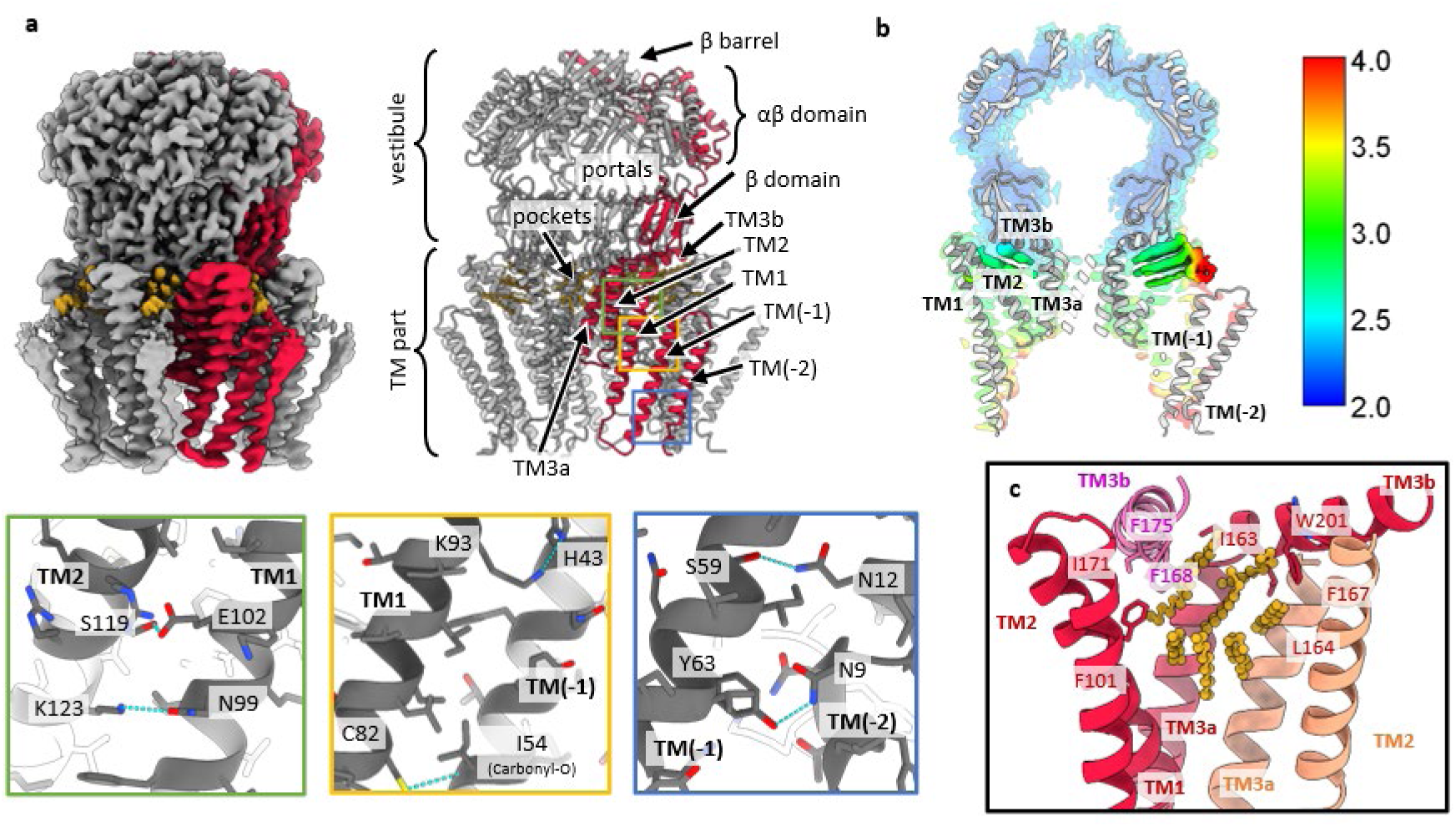
Structure of closed YnaI. **a** On the left, the cryo-EM map of closed YnaI acquired under condition I+ is shown in grey with one subunit highlighted in red and acyl chain densities in yellow. The right panel shows the corresponding model in the same orientation and color code, and labels of relevant structural parts and domains. Close-ups of the green, yellow, and blue boxes are shown below and show the intra-subunit hydrogen bonding network (blue dashed lines) between adjacent sensor paddle helices. **b** A slice of the model of closed YnaI in light grey is overlayed onto the corresponding slice of the map which is colored according to the local resolution of the map. The color key is shown on the right. **c** One pocket only is depicted, and the helices and residues are colored by subunit. One pocket is spanned by the helices TM1, TM2, TM3a, and TM3b from one subunit (red), TM2 and TM3a from one adjacent subunit (salmon), and TM3b from the adjacent subunit on the other side (pink). The eight modelled C_12_-alkyl chains are shown in yellow.

The map of closed YnaI resolves the transmembrane part of YnaI better than previous maps (figure 1a,b; figure S3) (Yu *et al*, 2018; Flegler *et al*, 2020; Hu *et al*, 2021; Catalano *et al*, 2021) suggesting structural stabilisation. We attribute this stabilisation to the added lipids similar as observed previously for MscS (Flegler *et al*, 2021). In the lipid supplemented YnaI we could model eight acyl chain densities per subunit in the cytosolic pockets, yet we could not distinguish whether they were contributed by detergent or lipid. The eight acyl chains formed parallel bundles with relative orientations that were compatible with four phospholipid molecules in a membrane context (figure 1b,c). The bundles were wedged between three adjacent subunits. The central of these three subunits generated most of the pocket with contributions from TM1, TM2 and TM3a/b. The other two subunits contributed either TM2 and TM3a or TM3b (figure 1c). TM3b formed van der Waals interactions with acyl bundles in two adjacent pockets. While its Phe168 and Phe175 coordinated the acyl chain in one pocket, Phe167 contacted the acyl chain in the next bundle. Thus, the dispersed acyl chain bundles were part of a hydrophobic inter-subunit interaction network that was located between the transmembrane part and the vestibule outside of the plane of the membrane. The pockets in the cytosol were continuous with the grooves between the paddles implying that lipids can enter the pockets from the reservoir of the membrane.

#### Opening YnaI

To generate an open conformation of YnaI, we introduced the mutation A155V (YnaI^A155V^). This mutation is analogous to A106V in MscS (Wang *et al*, 2008), A320V in MSL1 (*A. thaliana*) and G924S in MscK (*E. coli*) (Deng *et al*, 2020; Mount *et al*, 2022), which all show open pores in their respective structures. However, YnaI^A155V^ purified with standard DDM concentrations produced the closed state (figure S1, figure S4). Also the exchange of DDM to LMNG Lauryl Maltose Neopentyl Glycol (LMNG, condition II), which stabilises MscS most efficiently in the open state (Flegler *et al*, 2021) (figure S1, figure S2), did not result in open WT-YnaI or open YnaI^A155V^. However, higher concentrations of DDM throughout the purification, which is sufficient to open MscS (Flegler *et al*, 2021), also stabilised the majority of YnaI channels in the open state (figure S1, figure S2, figure S5). Open WT-YnaI channels with an overall resolution of 2.3 Å, and the overall structure of open YnaI^A155V^ were indistinguishable (figure S4). This implicated that WT-YnaI and YnaI^A155V^ adopted the same open conformation under the same purification conditions. The most prominent feature of open YnaI is the diminishing of the kink at Gly160, which separates the pore helices TM3a and 3b. Additionally, on the periplasmic side the helix TM3a kinks outwards at Gly149-Gly150, resulting in a wide-open pore (figure 2). Because Trp29 anchors the outer paddle helix TM(-2) to the cytosolic leaflet, the pockets in the open state are pulled into the plane of the cytosolic membrane leaflet and the pocket lipids align with the lipids of the cytosolic leaflet.

**Figure 2:**
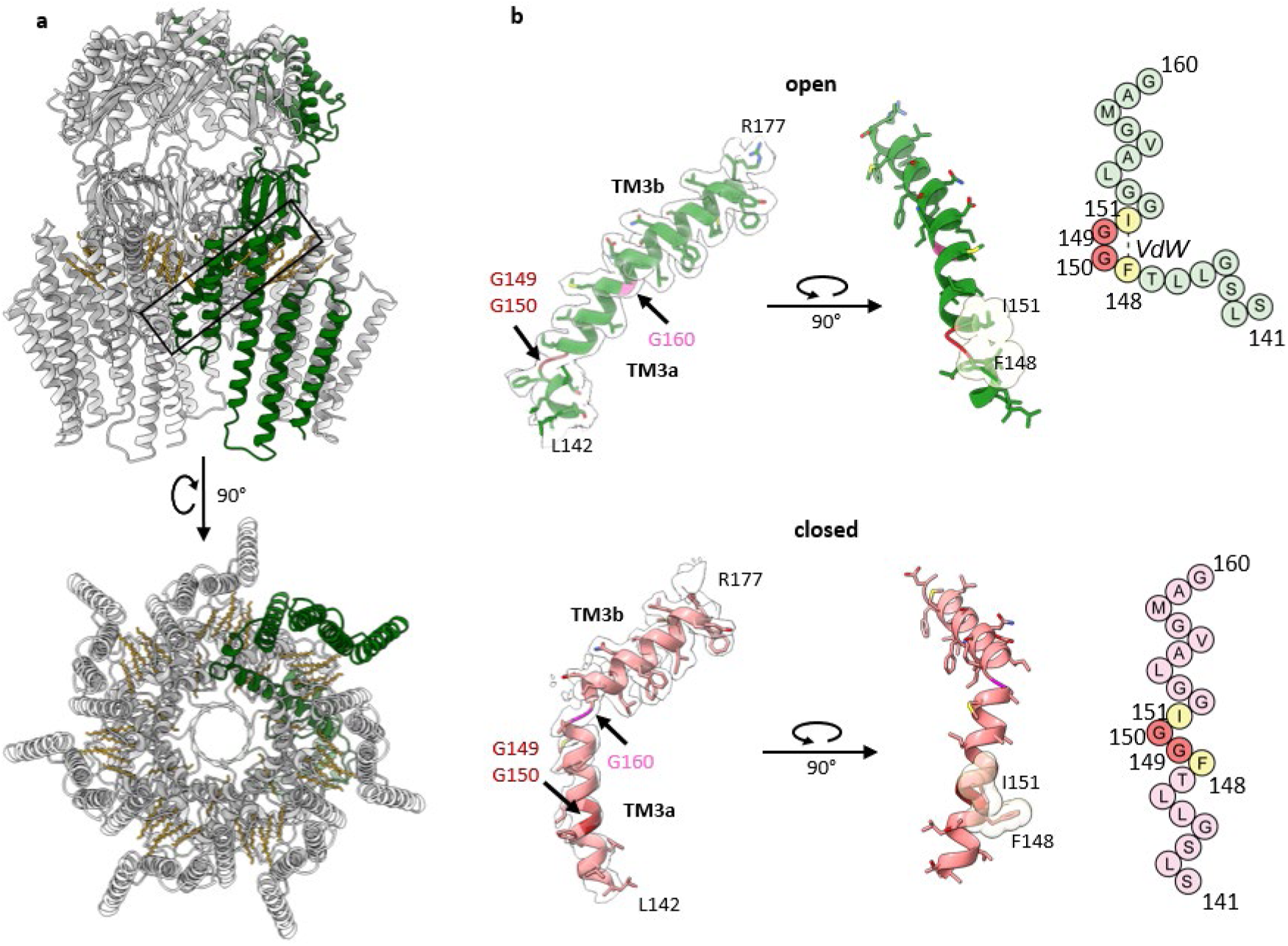
Structure of open YnaI. **a** The model of open WT-YnaI is shown in grey with one subunit in green. Alkyl chains are shown as yellow sticks. The pore helices TM3a-3b indicated by the black box are isolated shown in b. The model of open YnaI is viewed from the periplasmic side along the pore axis, showing a wide pore. **b** At the top, the conjoined helix is shown in its corresponding density. The kink at Gly160 (pink), which divides the pore helices TM3a and TM3b in the closed conformation, diminishes, while a novel kink at Gly149-Gly150 (red) is created in the open conformation. Gly149-Gly150 are flanked by two large hydrophobic residues, Phe148 and Ile151 which interact via Van der Waals bonds in the open state, stabilizing the kink. The van der Waals radii are shown as transparent yellow spheres. On the right, a scheme of the helix is depicted. At the bottom, the helices TM3a and TM3b are shown for the closed conformation analogously to the open conformation.

To assess whether the pore of the open WT-YnaI conformation supports free moving water as expected for an open conformation, we analysed the penetration pathway with GROMACS (Abraham *et al*, 2015) and found that the water phase is continuous between periplasm and vestibule with free moving water. Compared to closed YnaI (figure 3a), the free movement is enabled by increase in diameter from ∼9 Å to 22 Å in open YnaI and a rearrangement of the side chains making the interior of the pore less hydrophobic in the open state (figure 3b). Notable hydrophobic residues are Leu142 and Leu146, which render the periplasmic side of the closed pore hydrophobic but are moved outwards in the open state because of the outwards kinking of TM3a (figure 3c). Additionally, the electrostatic potential of the pore is unevenly distributed (figure 3d). MD simulations show preferential residence of cations and anions at different regions of the inner surface of YnaI in the open and closed state (figure S6a). In open YnaI, anions are attracted by Lys161 and reside with higher preference inside the open pore. In closed YnaI, Lys161 is no longer exposed, and the hydrophobic seal of Met158 and Leu154 expels ions from the pore but does not form a water-tight lock. The pore leads to an indentation at the periplasmic side, which is lined with negatively charged residues Glu66, Asp79, and Glu16 (figure 3e). The dense packing of charges in the closed state enriches cations in the indentation. This accumulation of ions agrees with a diffuse density at the periplasmic entrance of the pore that we observed in all maps of closed YnaI, but not in the maps of open YnaI (figure S1, figure S6b).

**Figure 3:**
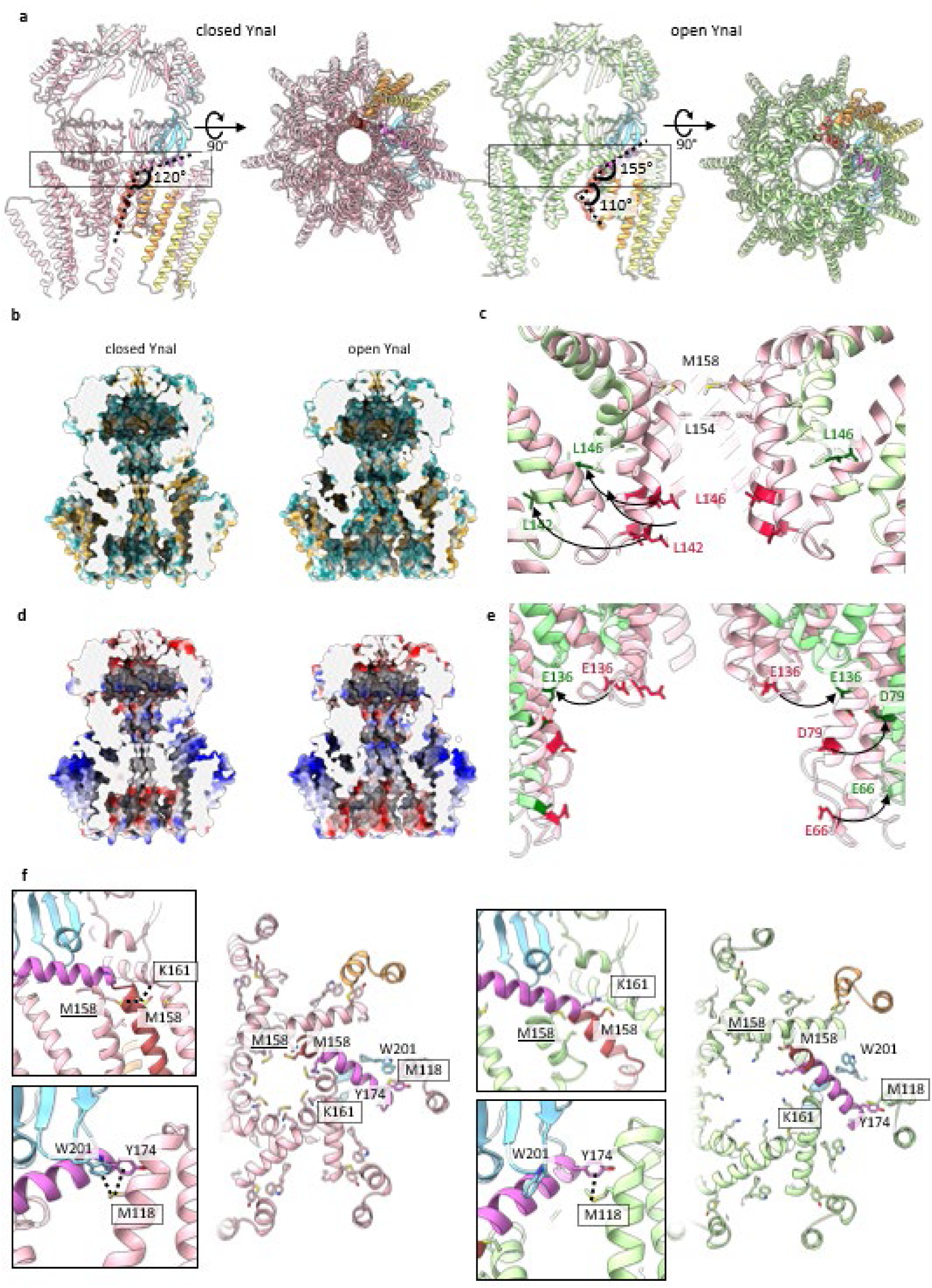
Comparison of closed and open YnaI. **a** Closed (light pink, left) and open (light green, right) are shown from the side and viewed along the pore from the periplasmic side. One subunit each is colored according to their organizational units: helices TM(-2) and TM(-1) are colored in yellow, and TM1 and TM2 in orange; the helix TM3a in red and TM3b in purple, the β- and αβ domain are depicted in light blue, and the β barrel in green. The dotted box indicates the position of the slice shown in f. **b** The surface of the models of closed (left) and open (right) YnaI is coloured by hydrophobicity. The front of the model is clipped for better visualisation of the pore. **c** Apart from the known hydrophobic constriction comprised of Met158 and Leu154, there are also Leu146 and Leu142 (red) in closed YnaI (pink) facing towards the symmetry axis. In the open conformation (light green) these leucines (green) protrude away from the channel axis, because the periplasmic part of the helix TM3a bends outwards. **d** The electrostatic potential of the surface is shown for closed and open YnaI. The front of the model is clipped for better visualisation of the pore. This highlights the accumulation of negatively charged residues at the periplasmic indentation. **e** The inner surface of the periplasmic indentation is highly electronegative, owing to the residues Glu66, Aps79, and Glu136 (red) which face towards the pore axis ion the closed conformation (pink). As the indentation becomes wider in the open conformation (light green), these residues (green) are spaced more and partly rotate away from the pore axis. **f** The color scheme is the same as in a. A slice is shown that encloses the cytoplasmic part of the pore with relevant residues labelled. Interactions are enlarged in the boxes. In the closed conformation, Met158 of one subunit can interact with the Met158 (underlined) of one adjacent subunit and Lys161 (boxed) of the other. Furthermore, a methionine aromatic bridge is present between Trp201 and Tyr174 of one subunit and Met118 (boxed) of a neighboring subunit. In the closed conformation, the Met158 is the diameter constricting residue of the pore, while it is K161 in the open conformation. In the open conformation, Met158 can interact with neither residue of an adjacent subunit because of the wide-open pore. Also, the methionine aromatic bridge is broken, because Trp201 tilts away from the other involved residues.

#### YnaI opening transition

The availability of the closed and the open conformation of YnaI allowed us a closer look at the underlying mechanism. Upon opening, the gross architecture of the cytosolic vestibule stays largely the same, while the diameter of the TM part increases on the periplasmic side (figure 3a). In the open state, the pore is wider and shorter than in the closed state – the effective pore length is reduced from ∼35 Å to ∼20 Å. The increase in the pore diameter is concomitant with tilting of the pore helices, rejoining TM3a/3b into a long continuous helix, while shortening TM3a towards the periplasm by kinking it outward. Joining of TM3a and TM3b changes the angle between the helical backbones at their pivot point G160 from 120° in the closed conformation to 155° in the open state showing that the newly formed helical segment is still bent (figure 3a). The straightening of TM3a/3b is reminiscent of the structural rearrangements of the pore helices of the large MscS-like channels MSL1 and MscK (Deng *et al*, 2020; Mount *et al*, 2022). At the periplasmic side the helical breakpoint in TM3a is located at Gly149 and Gly150, which are part of the ^149^GGIGG^153^ motif previously described (Flegler *et al*, 2020), and the chain continues as a short helix from Leu146 until Ser141 at the periplasmic side of the pore entrance. The angle spanned between the two helix portions is 110° (figure 2b, 3a). We noticed that kinking at Gly149 and Gly150 enables the two large flanking residues Phe148 and Ile151 to form van der Waals bonds that stabilise the kink in the open conformation (figure 2b). Kinking of TM3a moves the connected sensor paddle as an almost rigid body outward. In addition, the periplasmic ends of the helices tilt outwards by ∼5° toward the cytosolic side, which gives the TM region a less tapered appearance in the open state (figure 3a). The diameter of the periplasmic side of the TM part increases from 67 Å in the closed state to 90 Å in the open state while the diameter at the cytosolic side remains unchanged (figure S6c).

Next, we investigated the inter-subunit interactions that change between the open and closed conformation. In the closed but not open conformation the pore lining Met158 interacts with Met158 of one adjacent subunit and with Lys161 of the other adjacent subunit. Similarly, Trp201 in the β domain interacts with Met118 at the cytoplasmic end of helix TM2 of the adjacent subunit in the closed state (figure 3f) but pivots away in the open state. In the outer TM region, Asn8 is near Lys71 of the neighbouring subunit, and could form a H-bond with it (figure S6d) but loses the interaction in the open state, when the distance between these two residues increases to >15 Å. Comprehensively, upon opening most inter-subunit interactions in the TM region of YnaI are lost.

In the maps of the open channel, we resolved six acyl chains per pocket (figure 4a). We noticed that the arrangement of the acyl chains between the open and closed state differs: With respect to the membrane plane, the angles of the chains change from approximately 15° in the closed conformation to 35° in the open conformation. One of the acyl chain densities enters the vestibule close to Lys161 via the pockets in the open conformation. There it intercalates between two adjacent pore helices and stabilises the larger pore diameter. The gap for the acyl chain is generated by the straightening of helix TM3a-3b that retracts the Phe168 side chain in the helix TM3b, which would otherwise occlude the gap (figure 4b) consistent with the reported high lipid accessibility of Phe168 (Böttcher *et al*, 2015). Towards the plane of membrane, the cytosolic pocket is closed by a hydrophobic barrier formed by Leu41 and Ile124 of neighbouring subunits. This barrier is within the cytosolic leaflet of the membrane and blocks exchange of lipids between pockets and membrane in the open conformation but not in the closed conformation (figure 4c).

**Figure 4:**
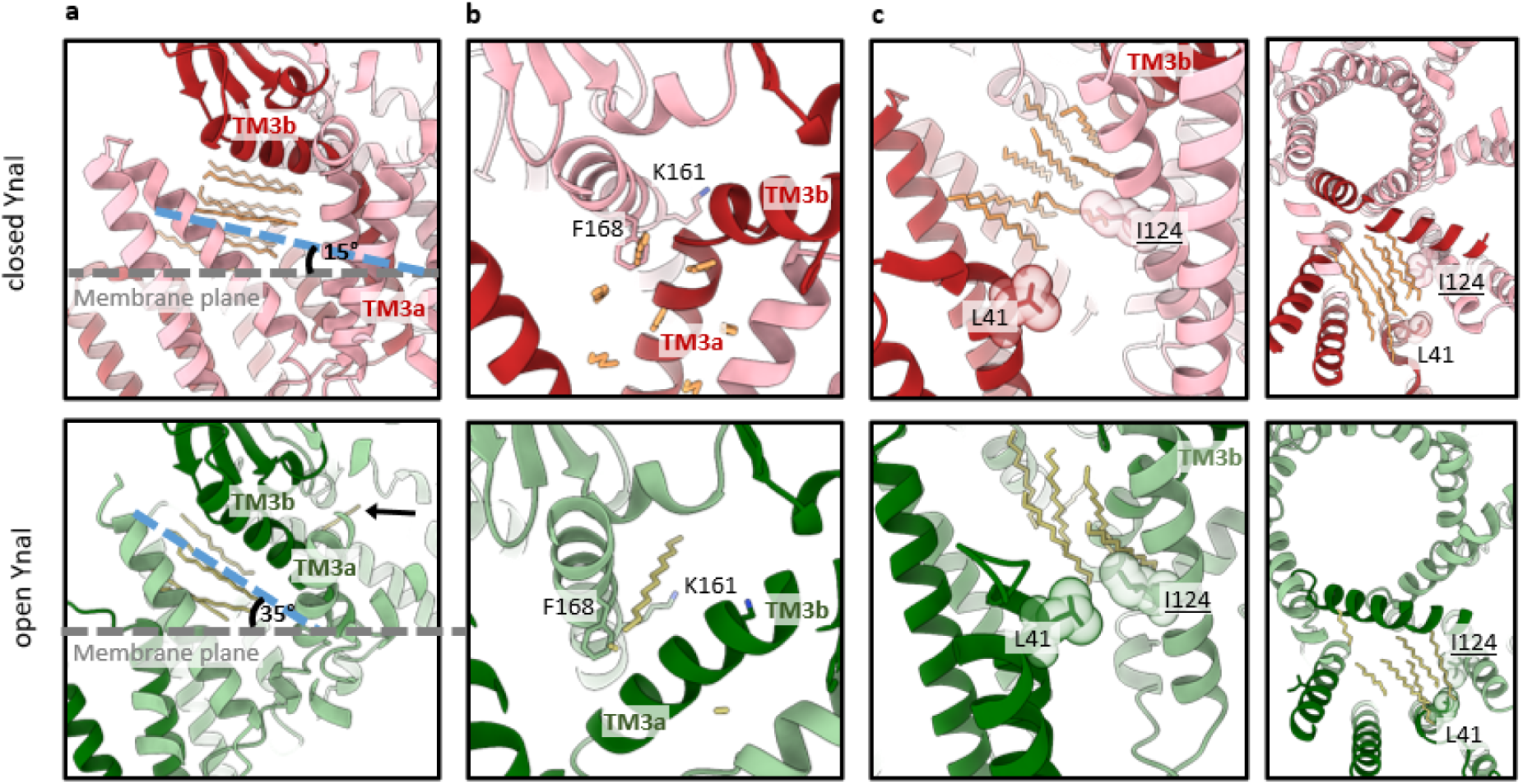
Reorganization of the alkyl chains in the pockets of closed and open YnaI. The figure shows the modelled alkyl chains as sticks of closed (top, red) and open (bottom, green) YnaI. One subunit each is highlighted in a darker colour. **a** The pocket lipids change their orientation as a bundle and with respect to the membrane plane (grey dotted line), the angle is increased from 15° to 35°, following the change of the helix TM3b. **b** Upon opening, a gab arises at Phe168 due to the straightening of helix TM3a-3b. In the open conformation, one alkyl chain enters this gap, showing a different tilt than the other pocket lipids. This alkyl chain is also indicated by the black arrow in a. **c** In the open state, a hydrophobic bridge is built between Leu41 of one subunit and Ile 124 of a neighboring subunit. Their van der Waals radii are depicted as transparent spheres. This bridge probably prevents the pocket lipids from leaving the channel in the open conformation and could play a role in closing the channel when membrane tension is released.

#### Open and closed conformations of a YnaI-MscS chimera

Our structural analyses of closed and open YnaI do not show molecular interactions between the paddle helices and the pore and vestibule. This suggests that the paddle (the sensing module) moves independently from the pore and vestibule (the conductivity module). To further investigate this hypothesis, we explored a chimeric construct comprising the sensing module of YnaI, and the conductivity module of MscS. As splitting point, we defined Gly139 of YnaI, or Gly90 of MscS, respectively. This glycine residue is conserved within the *E. coli* MscS-like family members (figure 5a). Electrophysiology showed that this new channel exhibits the properties of both YnaI and MscS: Its current is comparable to that of MscS (∼1 nS (chimera) vs. 1.2 nS (MscS)), while its pressure threshold for opening P_MscL_:P_YnaI-MscS_ is similar to YnaI (1.02 (chimera) vs. 1.04 (MscS)) (figure 5a,b). This indicates that specific channel properties are restricted to either the sensing or the conductivity module. However, in the chimera the ability of the channel to open stably and/or remain open seems to be disrupted: it does not show the typical staircase-like opening behaviour as wildtype channels, but opens mostly flickery, and only a subset opens with very short dwell times (figure 5b). The chimera provides protection in hypoosmotic downshock assays, despite being only weakly overexpressed (figure S7a,b). The structure of the chimera purified under condition I+ represents a closed state that shows the MscS-derived conductivity module unchanged (figure S8, figure 5c,d, figure S9a,b). The YnaI sensor paddles have still the same structure as observed for WT-YnaI but are slightly shifted and tilted as a rigid body compared to it (figure 5 c,d,e). As a result, the innermost paddle helix TM2 aligns with that of WT-MscS. We also observed the designated pore lipids of closed MscS (Reddy *et al*, 2019; Zhang *et al*, 2021), indicating that accumulation of lipids is a property of the MscS pore probably related to its high hydrophobicity (figure S9c,g). The helical backbones of the pore helices are highly similar in the chimera, YnaI, and MscS (figure 5d). The chimera purified under condition III shows an open state with a novel pore architecture (figure S8, figure 5c,d, figure S9e,f,g). Upon opening, the angle between helices TM3a and TM3b decreases from 135° to 100°, which is more than in WT-MscS (135° to 120°). This leads to a funnel-like shape of the chimera pore and results in a unique positioning of the sensor paddles compared to the conductivity module, while they remain a rigid body (figure 5c,d). The outward movement of helix TM3a shifts them further clockwise (viewed from the periplasmic side) than in WT-MscS, but the paddles do not tilt within the plane of the membrane. The studies on the chimera exemplify that the sensing and conductivity modules function independently but adapt in their conformational response to the overall channel organisation. The chimera showed that two modules derived from different channels resulted in a functional phenotype that reflects the features of the original channels, meaning that these sensing and conductivity modules are interchangeable. Yet, the artificial channel also shows unique characteristics, like the unstable or short openings. Conclusively, we learned from this specific chimera 1.) that the behaviour of the kink dividing the helices TM3a and TM3b is a property inherent to these helices, 2.) that Gly149-Gly150 of YnaI are indeed necessary to kink the helix TM3a outwards, and 3.) that tilting of the paddles within the membrane plane is not related to the pore.

**Figure 5:**
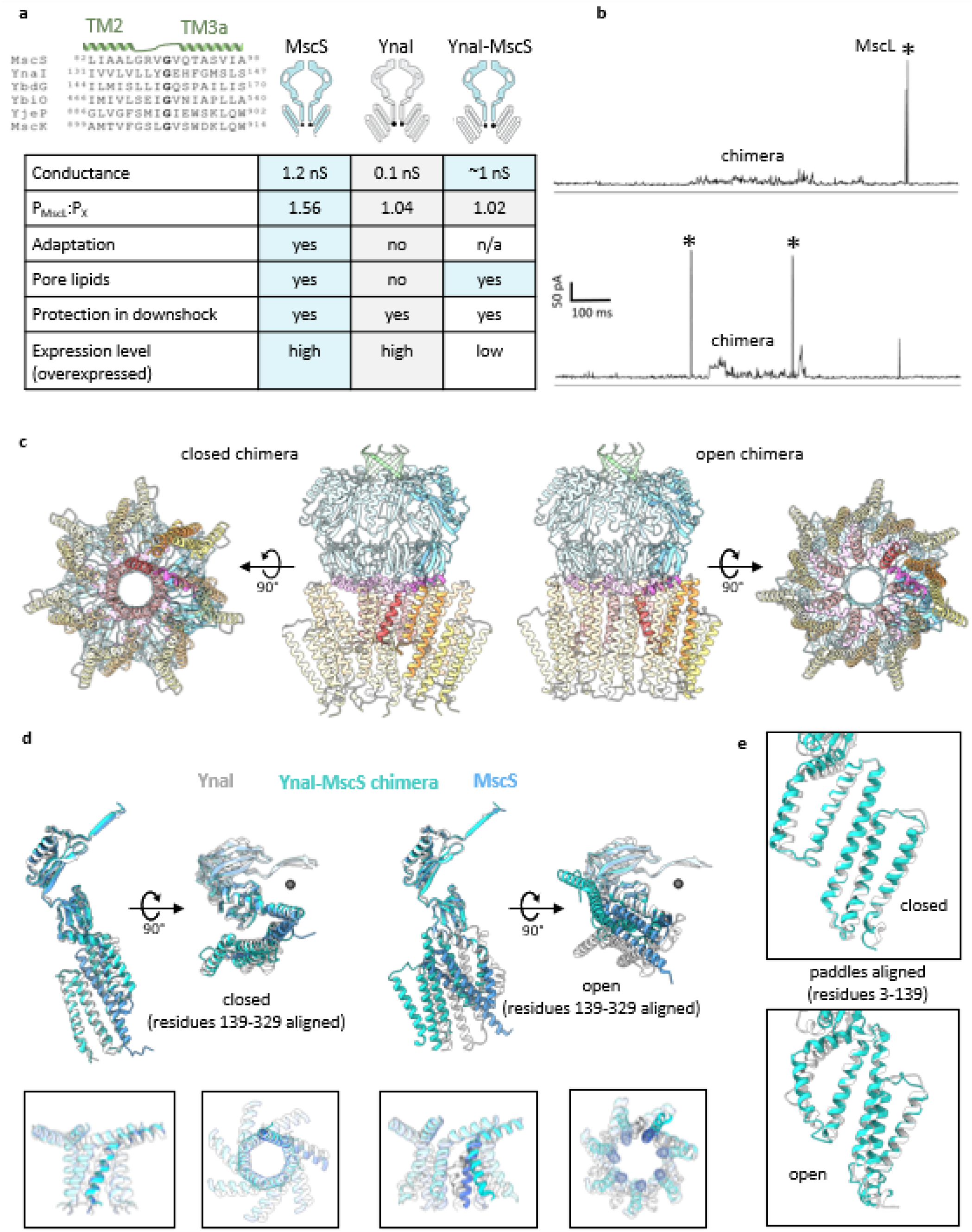
**a** The YnaI-MscS chimera was designed based on a glycine (bold) in the loop connecting the helices TM2 and TM3a. This glycine is highly conserved within the *E. coli* MscS channel family, which is Gly139 in YnaI (grey in right scheme), and Gly90 in MscS (blue in right scheme), respectively. The YnaI-MscS chimera shows characteristics of both donor channels, YnaI and MscS, but also novel features. **b** Representative current traces from membrane patches from spheroplasts from *E. coli* MJF429 (ΔMscK, ΔMscS) expressing the YnaI-MscS chimera are shown with MscL labelled with an (*). The chimera shows both a flickery phenotype and/or short openings. **c** The models based on cryo-EM maps are shown, colored according to their organizational units: helices TM(-2) and TM(-1) are colored in yellow, and TM1 and TM2 in orange; the helix TM3a in red and TM3b in purple, the β- and αβ domain are depicted in light blue, and the β barrel in green. One subunit is darker. While the architecture of the closed chimera largely resembles that of YnaI, it shows a novel pore arrangement in the open conformation. The helices TM3a are slightly bent outwards at the periplasmic side, resulting in a funnel shaped pore. Additionally, the paddle helices rotate clockwise in the open form – a feature that is observed for open MscS. The four paddle helices of one subunit remain a rigid body. **d** The pore and cytosolic domains of the closed structures are aligned. YnaI is shown in white, MscS in blue, and the chimera in cyan. One subunit is shown from the side (left) and the top (right). The black sphere marks the symmetry axis. The inserts in the black boxes represent the aligned pores (TM3a-TM3b) showing all seven subunits. The pore and cytosolic domains of the open structures are aligned. c The pore and cytosolic domains of the open structures are aligned. One subunit is shown from the side (left) and the top (right). The black sphere marks the symmetry axis. The inserts in the black boxes represent the aligned pores (TM3a-TM3b) showing all seven subunits. **e** The paddles (residues 3-139) of the closed chimera and YnaI (top) and open chimera and YnaI (bottom) are aligned showing that the paddle organization is essentially unchanged.

## Discussion

Knowledge about different conformational states of the MscS-like channels is inevitable for understanding the underlying molecular basis of mechanosensation. Our new high-resolution structures of the larger MscS paralogue YnaI and a chimeric channel of YnaI and MscS allows the identification of shared and unique features of pore opening among MscS-like channels. A structural hallmark of opening MS channels based on the currently available structures is an area expansion of the transmembrane domain (Bavi *et al*, 2023; Mount *et al*, 2022; Deng *et al*, 2020; Zhang *et al*, 2021; Jojoa-Cruz *et al*, 2022; Flegler *et al*, 2021). The force-from-lipids principle states that the anisotropic forces of the lipid bilayer (Cantor, 1998, 1997) and their changes directly enforce channel gating (Martinac *et al*, 1990; Kung, 2005; Teng *et al*, 2015). In a membrane under tension, the channel conformation with a larger membrane cross section is energetically more favourable. Hence, a simple in-plane area expansion can dispense the energy for conformational changes (Wiggins & Phillips, 2005). This can be described with:

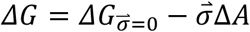

Where 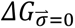 is the free energy difference for channel opening without tension applied, *σ⃑* is the membrane tension, and Δ*A* the in-plane area change of the membrane cross section upon opening (Haswell *et al*, 2011; Guo & MacKinnon, 2017). Furthermore, the sensitivity of an MS channel towards tension is given by:

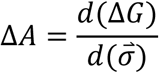

This means that a higher area expansion upon gating at the same tension infers a higher mechanosensitivity (Haswell *et al*, 2011; Guo & MacKinnon, 2017). We noticed for YnaI, that an area expansion only takes place at the periplasmic side of the TM part, while the area at the cytosolic side remains almost the same. The same is true for MscS opening. But while the periplasmic in-plane area expansion for MscS is 22 nm^2^, it is only 14 nm^2^ for YnaI. YnaI opens at significantly higher membrane tension levels than MscS (Edwards *et al*, 2012), which fits the observed different in-plane area expansions for MscS and YnaI.

Although area expansion seems to be the common basis for gating mechanosensitive MscS-like channels, there are further aspects to be considered. We could model numerous acyl chains in our maps of closed and open YnaI. Though the origin of these acyl chains could not be clearly attributed to either detergent or lipid molecules, their differing positions and orientations between the closed and open conformation are informative. This is in agreement with observations by (Park *et al*, 2023) based on coarse-grained MD simulations. These simulations indicated a re-orientation of the pocket lipid molecules of MscS and MSL1, stating that lipid reorganisation and not lipid removal accompany channel opening. Park *et al*. stated that membrane tension also favours an open channel conformation because the membrane perturbation in the pockets of the closed channel is energetically less favourable.

Our studies on a YnaI-MscS chimera indicated lipid coordination is an inherent feature and dependent on the molecular details of a channel. Specifically, we have observed that the presence of pore lipids is associated with the pore of MscS but not with the pore of YnaI which has a less hydrophobic interior.

Our open structure of YnaI shows the outwards bending of the pore helices TM3a at Gly149-Gly150, which divides TM3a into two sub-helices in the open conformation. Outward bending of the pore helices TM3a in YnaI is, together with diminishment of the kink between the helices TM3a and TM3b, the major contributor to the significant pore shortening of open YnaI to approximately 20 Å. A shorter pore length means a reduction of the transmembrane diffusion distance and a lowering of the intrapore diffusion energy barrier, reminiscent of K^+^ channels which only have a relevant pore length of 12 Å (Zuo *et al*, 2024; Esfandiar *et al*, 2017; Zhou *et al*, 2020; Doyle *et al*, 1998). Nonetheless, it is curious why YnaI opens to such a short pore with a large width, while the conductance is limited by the narrow lateral portals in the vestibule (Flegler *et al*, 2020; Hu *et al*, 2021). This suggests that in the open channel ions can easily exchange between vestibule and periplasm while entering the vestibule from the cytosol remains restricted.

The chimera cannot reduce the kink dividing the helices TM3a and TM3b and cannot bend the helix TM3a outwards. Because of the missing di-glycine hinge, the paddles rotate clockwise similarly as in MscS. Instead, pore shortening in the chimera is realised by reducing the kink angle which give rise to the observed funnel-shaped pore and is also accompanied by area expansion on the periplasmic side.

Comprehensively, our present study describes numerous structural features of YnaI, which are either unique or shared with other MscS-like channels. Lipids were postulated to initiate and control MscS gating (Flegler *et al*, 2021; Zhang *et al*, 2021; Pliotas *et al*, 2015). The new findings indicate that pocket lipid removal cannot initiate gating in YnaI, as indicated by the fully crowded pocket, and highlight that area expansion following membrane tension is the major driving force for rearranging the protein scaffold towards a wider pore. Even more, we attribute a novel function to lipids, speculating that pocket lipid orientation might provide a mechanism for pore closing: lipids might push against the bridge and become continuous with the pocket lipids, resulting in a reorientation of the pocket lipids. The lipid tails press against the pore and enforce kinking of the pore helices and thereby pore closure. Based on our high resolution YnaI maps, which allowed reliable model building of most of the closed and open channel, we suggest the following model for gating (figure 6): A tensed membrane enforces an area increase and shape change of the TM part because an expanded conformation is then energetically supported. Area expansion on the periplasmic side imposes an outward translocation of the sensor paddles which induces bending of the connected pore helix TM3a at Gly149-Gly150. Concomitantly, the angle between TM3a and TM3b at Gly160 increases, leading to a decrease of the kink separating these helices, and as a result, the pore is opened. In YnaI, the pocket lipids are separated from the bilayer lipids by a hydrophobic bridge and can therefore not exchange with those or leave the pockets in the open state. The rearranged open pore helices enforce a reorientation of the lipids and the radial area that is occupied by lipids decreases upon pore opening. This is necessary, because the diameter of the cytosolic side of the TM part merely increases upon opening. YnaI highlights that although tension-induced area expansion is a common feature of mechanosensitive ion channels there are certain features for fine-tuning their function.

**Figure 6:**
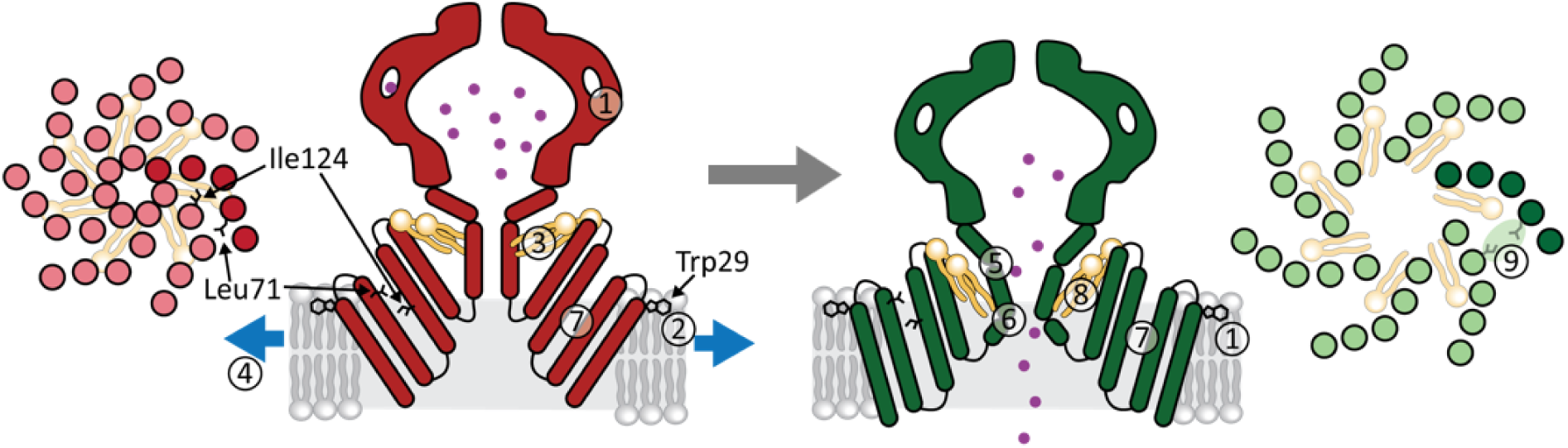
The gating transition of YnaI. YnaI is schematically shown in its closed (red), and open (green) conformation. Two subunits are shown for the side views, and the TM helices of all seven subunits viewed from the periplasmic side are shown at the very left and very right, respectively. In these top views, one subunit is colored in a darker colour. In a model for gating YnaI, several aspects have to be considered. The portals (1) in YnaI are limiting for the conductance. The Trp29 (2) anchors the TM domain at the interface of the cytosolic membrane leaflet. In closed YnaI, lipids (3) fill the hydrophobic pockets. These lipids lie almost parallel to the membrane plane. Tension (4) leads to a then energetically favoured area increase of the periplasmic side of the TM part. The kink dividing the helices TM3a and TM3b diminishes (5), and the helix TM3a bends outwards (6), resulting in shortening of the pore. The paddles are relocated as a rigid body (7), and concomitantly, the pocket lipids (8) change their orientation to be more perpendicular with respect to the membrane plane. Radial relocation of the paddles enables the residues Leu71 and Ile124 to form a hydrophobic bridge in the open conformation (9), which separates the lipids in the pockets from the cytosolic membrane leaflet and reducing exchange with the lipid bilayer in the open state. In the open state, the squeezed pocket lipids press against the pore, ultimately enforcing its closure.

## Methods

### Mutations

The pTrc_YnaI-LEH_6_ construct (Edwards *et al*, 2012) was used as a template to introduce the TEV cleavage site, ENLYFQS by use of the Q5 site-directed mutagenesis kit (NEB). The YnaI^A155V^ mutant was generated based on pTrc_YnaI-ENLYFQS-LEH_6_ following the Stratagene QuikChange protocol using Q5 polymerase (NEB). Mutations were confirmed by sequencing (Microsynth Seqlab). The YnaI-MscS chimera synthetic construct with a C terminal His_6_ tag was purchased from Thermo Fisher Scientific and was subsequently cloned into a pTrc vector. The TEV cleavage site ENLYFQS was introduced using the Q5 site-directed mutagenesis kit (NEB).

### Expression and purification of YnaI constructs

Expression and purifications of the different YnaI constructs and applied purification approaches have been described previously for MscS (Flegler *et al*, 2021) with minor modifications. MJF641 cells (“Δ7”) (Edwards et al, 2012) that overexpressed YnaI constructs with a C-terminal His_6_-tag were resuspended and homogenised in solubilisation buffer containing either 1 % (w/v) DDM (Glycon) (condition I and I+) or 1.5 % (w/v) DDM (condition III) incubated for 1 h on ice. After centrifugation, YnaI was filtered and applied onto prepacked 0.5 ml Ni-NTA agarose columns and washed with 40 ml washing buffer containing either 0.05 % DDM (condition I and I+) or 0.5 % DDM (condition III), 50 mM sodium phosphate buffer pH 7.5, 300 mM NaCl, 10% (w/v) glycerol, and 20 mM imidazole. For YnaI in LMNG, the washing buffer contained 0.03 % LMNG instead of DDM (condition II). For elution, the same buffer was used but with 300 mM imidazole. The protein containing elution fraction was further purified on a Superose 6 10/300 size exclusion column (GE Healthcare) equilibrated with SEC buffer (50 mM HEPES pH 7.5, 150 mM NaCl, and 0.03 % DDM or 0.02 % LMNG, respectively. For closed YnaI with lipid supplementation (condition I+), the washing buffer contained 0.7 mg/ml Azolectin (Sigma) and the SEC buffer 0.5 mg/ml Azolectin. Peak fractions were concentrated using centrifugal filter units with a MW cutoff of 100 kDa (Amicon) to 4-6 mg/ml.

### Cryo-EM

Quantifoil 300 mesh R 1.2/1.3 copper grids were glow-discharged in air for 2 min at medium power in a PDC-002 plasma cleaner (Harrick). Subsequently, 3.5 μl sample were applied on the grids and plunge frozen in liquid ethane using a Vitrobot IV (FEI/ Thermo Fisher) with following parameters: wait time 0 s, drain time 0 s, blot time 5 s, blot force +25, chamber humidity set to 100 % and chamber temperature set to 4 °C.

For data acquisition, vitrified grids were transferred to a Thermo Fisher Titan Krios G3 transmission electron microscope. For the data sets of the closed and open WT-YnaI condition I+ and III), and the closed and open YnaI-MscS chimera, movies were acquired at 300 kV with a Falcon 4i detector and a Selectris energy filter with a slit width of 5 eV at a nominal magnification of 130 kx, which corresponds to a calibrated pixel size of 0.946 Å. A total dose of approximately 40-70 e^-^/Å^2^ (Table S1a and S1b) was used and data was collected with the EPU-acquisition software (Thermo Fisher). For the data sets of YnaIA155V (condition II and III and WT-YnaI in LMNG (condition II) movies were acquired at 300 kV with a Falcon 3 camera in integrating mode at a nominal magnification of 75kx, which corresponds to a calibrated pixel size of 1.0635 Å. A total dose of 80-90 e^-^/Å^2^ was used.

### Single particle analysis

The cryo-EM structures were solved with cryoSPARC versions 4.1-4.4 (Punjani *et al*, 2017). Movies were pre-processed in a cryoSPARC live session using patch motion and patch ctf for motion correction and dose weighting, and CTF estimation, respectively. Particles were picked using the blob picker with a diameter of 124 Å and extracted with a box size of 256 px, followed by two to three rounds of 2D classification to remove junk particles. As initial volume for refinement, an ab-initio reconstruction with C1 symmetry was generated for the initial refinements of the first WT-YnaI (condition I+). For other samples, previously obtained maps of YnaI were then used as initial models. The initial lowpass filter applied to the starting model was 30 Å. A heterogenous refinement with four classes and applied C7 symmetry was conducted to distinguish between different populations. This step proved essential to differentiate between the closed and open conformations. Different classes from the heterogeneous refinement were subjected to individual non-uniform refinements with C7 symmetry and enabled global CTF refinement. For open WT-YnaI, a 3D variability analysis of the best class from the heterogenous refinement was done, and the particles of the two best performing clusters were joined for a final non-uniform refinement. Maps were sharpened with an applied B factor of -10 to -30. The remaining aligned particles, and the final map, of open WT-YnaI, and the closed and open YnaI-MscS chimera were additionally transferred from cryoSPARC to Relion 5 (Scheres, 2012; Kimanius *et al*, 2024) using pyem (Asarnow *et al*). In Relion, a 3D classification without blush regularisation was performed with the C7 symmetry relaxed. The best class(es) were 3D refined with blush regularisation and relaxed C7 symmetry. Details for the individual data sets are given in Table S1a for closed and open YnaI and in Table S1b for the closed and open YnaI-MscS chimera.

### Model Building and Refinement

All model building was performed in *Coot* (Emsley *et al*, 2010) version 0.9.7, using existing structures as starting points (PDB 6ZYD (YnaI (Flegler *et al*, 2020)), PDB 6RLD (MscS (Rasmussen *et al*, 2019)). For most residues of the outermost helix TM(-2) of open YnaI, no clear side chains were visible, hence, the equivalent helices of the model of closed YnaI were placed in the density and fitted as a rigid body. For the open form of the YnaI-MscS chimera, no side chain densities were clearly attributable for the helices TM(-1) and TM(-2), therefore, also the helices were placed in the corresponding densities as rigid bodies. The nine C-terminal residues were resolved in neither of the maps. *Phenix* (Adams *et al*, 2010) version 1.17.1 was used for real-space refinement imposing secondary structure restraints and subsequently, the models were validated with *MolProbity* (Chen *et al*, 2010). Details for the individual data sets are given in Table S1. All map- and model images were created with *UCSF Chimera* versions 1.15 (Pettersen *et al*, 2004) or *UCSF ChimeraX* version 1.7.1 (Goddard *et al*, 2018; Pettersen *et al*, 2021).

### Electrophysiology

Giant *E. coli* spheroplasts were generated as described previously (Blount *et al*, 1999). Patch-clamp recordings were conducted on membrane patches derived from giant protoplasts using the strains MJF429 (ΔmscS, ΔmscK), and MJF641 (ΔmscS, ΔmscK, ΔynaI, ΔybdG, ΔybiO, ΔyjeP, ΔmscL) transformed with plasmids for overexpressing the YnaI-MscS chimera (Edwards *et al*, 2012; Levina *et al*, 1999). The cultures were induced with 1 mM IPTG for 2 h before protoplast generation. Excised inside-out patches were measured at +40 mV with identical pipette and bath solutions (200 mM KCl, 90 mM MgCl_2_, 10 mM CaCl_2_, 5 mM HEPES buffer at pH 7.0). Data was acquired at 3-kHz filtration using a HEKA EPC-8 amplifier and the Patchmaster software (HEKA). The pressure threshold for activation of the YnaI-MscS chimeric channels was referenced against the activation threshold of MscL (P_MscL_:P_YnaI-MscS_) to determine the pressure ratio for gating (Blount *et al*, 1996). Reference measurements resulted in P_MscL_:P_YnaI-MscS_ = 1.02 ± 0.05 (n = 8; derived from two spheroplast preparations from two different transformations).

### Whole-cell Western blot analysis

The expression level of the YnaI-MscS chimera was compared to that of MscS and YnaI by a whole-cell Western blot as described in the literature (Miller *et al*, 2003a, 2003b). Shortly, *E. coli* MJF641 cells were transformed with the target constructs, single colonies were purified, and overnight cultures were grown at 37 °C. The next day, precultures were inoculated with the overnight cultures, grown to an OD_600_ of 0.4, and a portion thereof was added to 30 ml of pre-warmed LB medium to give a starting OD_600_ of 0.05. The OD_600_ was monitored until 0.4, and cells were induced for 30 min with 0.3 mM IPTG. Subsequently, the OD_600_ was measured (OD_600,harvest_), and 1 ml of the cells were harvested by centrifugation (10000×g, 1 min), and the resulting pellet was resuspended in 1 ml of PBS buffer. For sample preparation for Western blotting, [2.2/OD_600,harvest_*100] μl of the harvested cells were centrifuged at 10000×g for 1 min, and the pellet was resuspended in 20 μl of solubilisation buffer (20 % glycerol, 2 % SDS, 0.12 M Tris-HCl (pH 6.8), 0.5 g/l bromophenol blue, 2 % (v/v) β mercaptoethanol). Samples were heated at 100 °C for 10 min before loading onto an SDS gel.

## Data availability

The EM maps of C7 processed closed and open YnaI (purified under condition I+ and condition III), and of the closed and open YnaI-MscS chimera (purified under condition I+ and condition III) obtained in cryoSPARC have been deposited in the Electron Microscopy Data Bank (EMDB) and corresponding atomic models have been deposited in the Protein Data Bank under following accession codes PDB 9H95 and EMD-51953 (closed YnaI), PDB 9H2P and EMD-51813 (open YnaI), PDB 9H2S and EMD-51817 (closed chimera), and PDB 9H2V and EMD-51818 (open chimera). Maps processed in Relion with C7-relaxed symmetry of open YnaI, and the closed and open YnaI-MscS chimera have been deposited to the EMDB under the accession codes EMD-51897 (open YnaI), EMD-51898 (closed chimera), and EMD-51954 (open chimera).

## Acknowledgements

We thank Christian Kraft for technical assistance. Electron Cryo Microscopy was carried out in the cryo EM-facility of the Julius-Maximilians-Universität Würzburg.

## Author Contributions

V.J.F. performed biochemical preparations, V.J.F., T.R. and B.B. analysed cryo-EM data. V.J.F. and A.R. conducted electrophysiology experiments. B.B. conceived funding and supervised the work. V.F. prepared the manuscript with contributions from all authors. All authors edited the manuscript.

---

**Figure S1:**
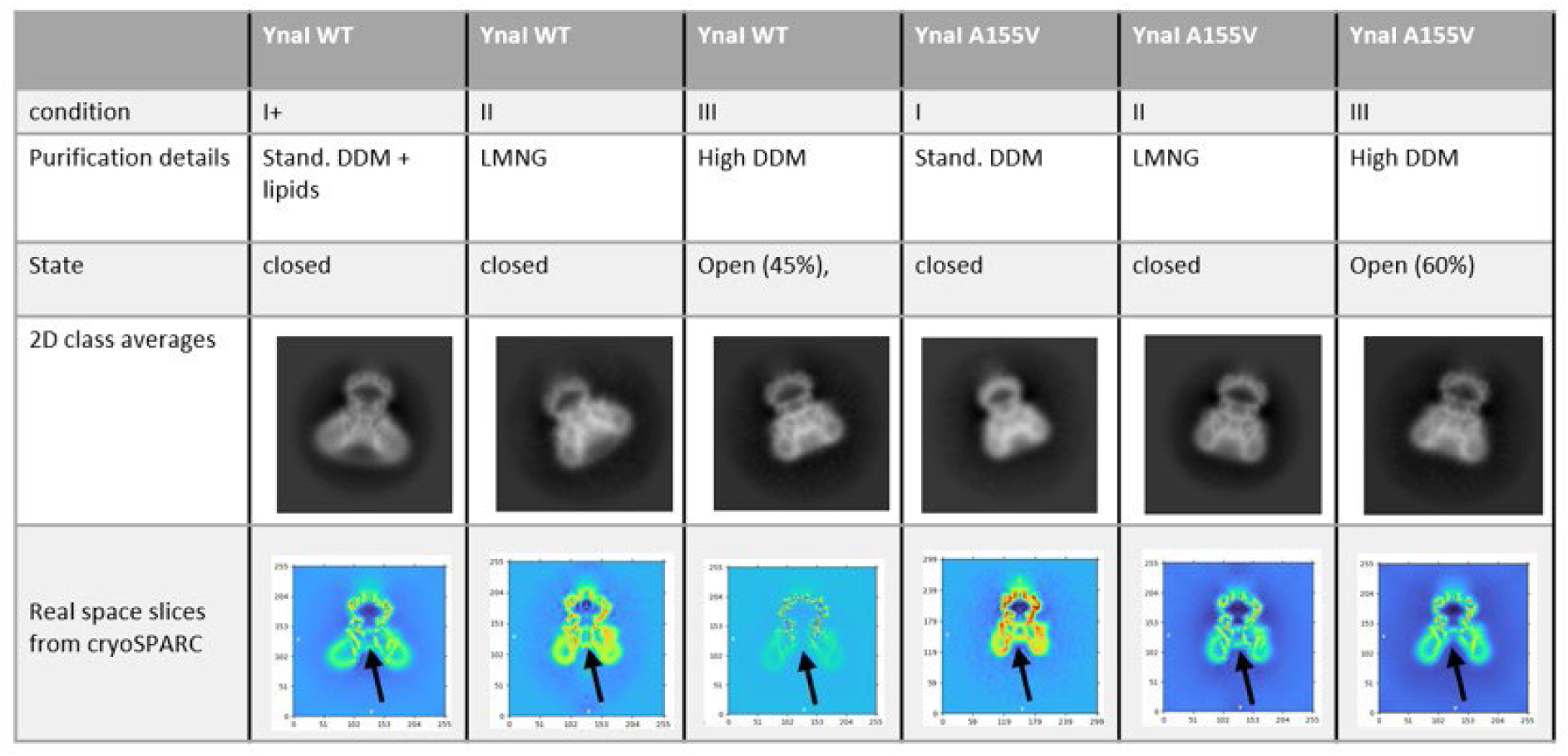
Summary of the different purification approaches. The table describes the different purification strategies applied to wt-YnaI and YnaI^A155V^ and the observed state of the purified protein after cryo-EM analysis. Furthermore, representative 2D class averages and map projections from the cryoSPARC output are shown. On a 2D class average level, the closed and open conformation are not distinguishable, but the map slice (taken from cryoSPARC) of the open conformation lacks the second density bridge within the periplasmic entrance of the pore (black arrows).

**Figure S2:**
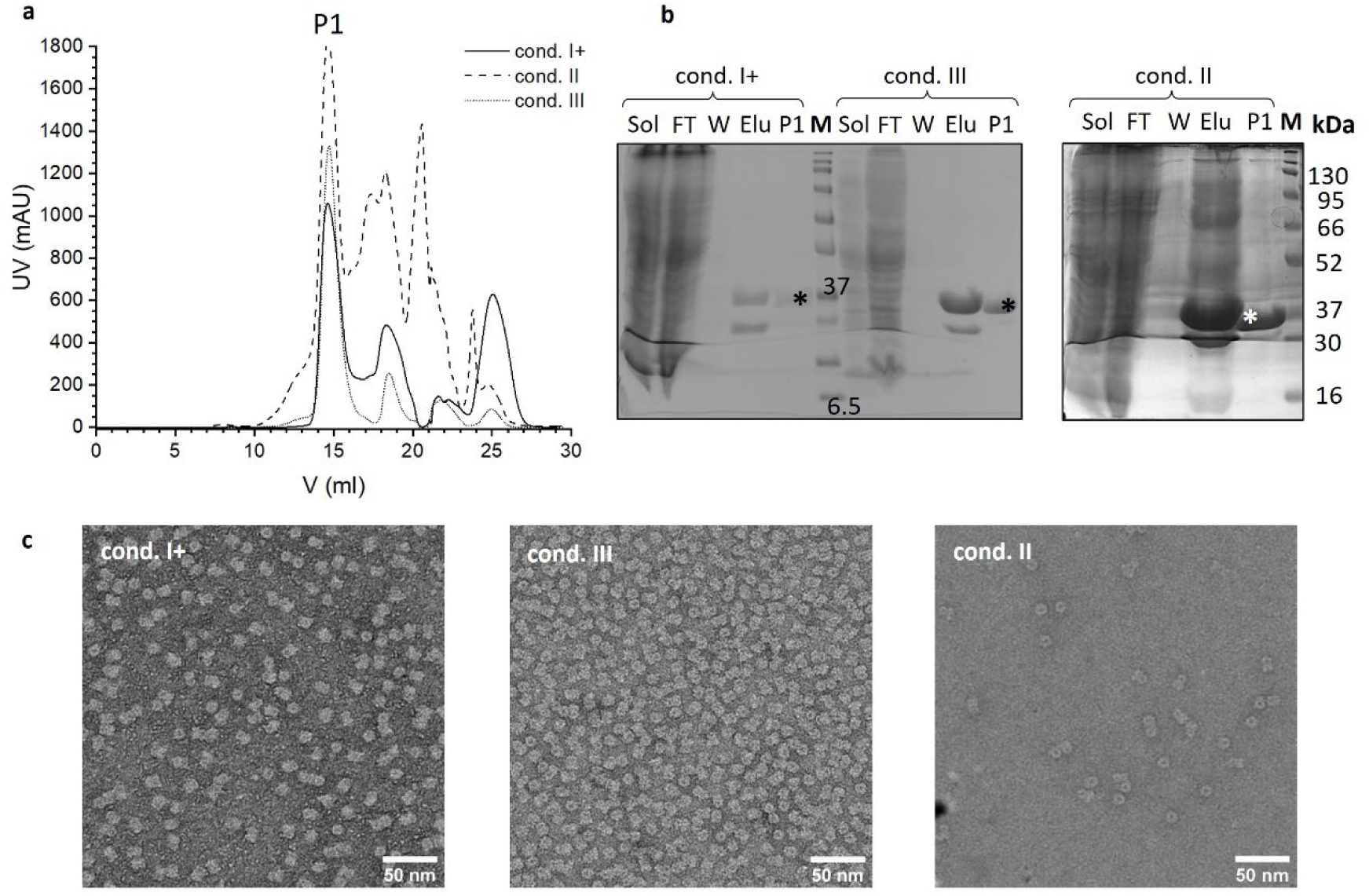
Purifications of WT-YnaI under the different conditions. WT-YnaI was purified under different conditions. **a** The size exclusion chromatograms are shown for YnaI purified under condition I+ (solid line), II (dashed line), and III (dotted line). The peak at 15 ml (P1) is the desired heptamer. **b** SDS-PAGE are shown for all three purification approaches, samples were taken from the supernatant after solubilisation (Sol), IMAC flowthrough (FL), IMAC wash (W), IMAC protein-containing elution fraction (Elu), and HPLC heptamer peak (P1), as well as a Marker (M). The signals from YnaI at the height of the 37 kDa marker signal are highlighted with an asterisk. **c** The P1 fraction was pooled and concentrated to the final concentration needed for subsequent cryo-EM preparation. Before, a dilution (1:2000-1:3000) of the concentrated sample was negatively stained and imaged for quality assessment.

**Figure S3:**
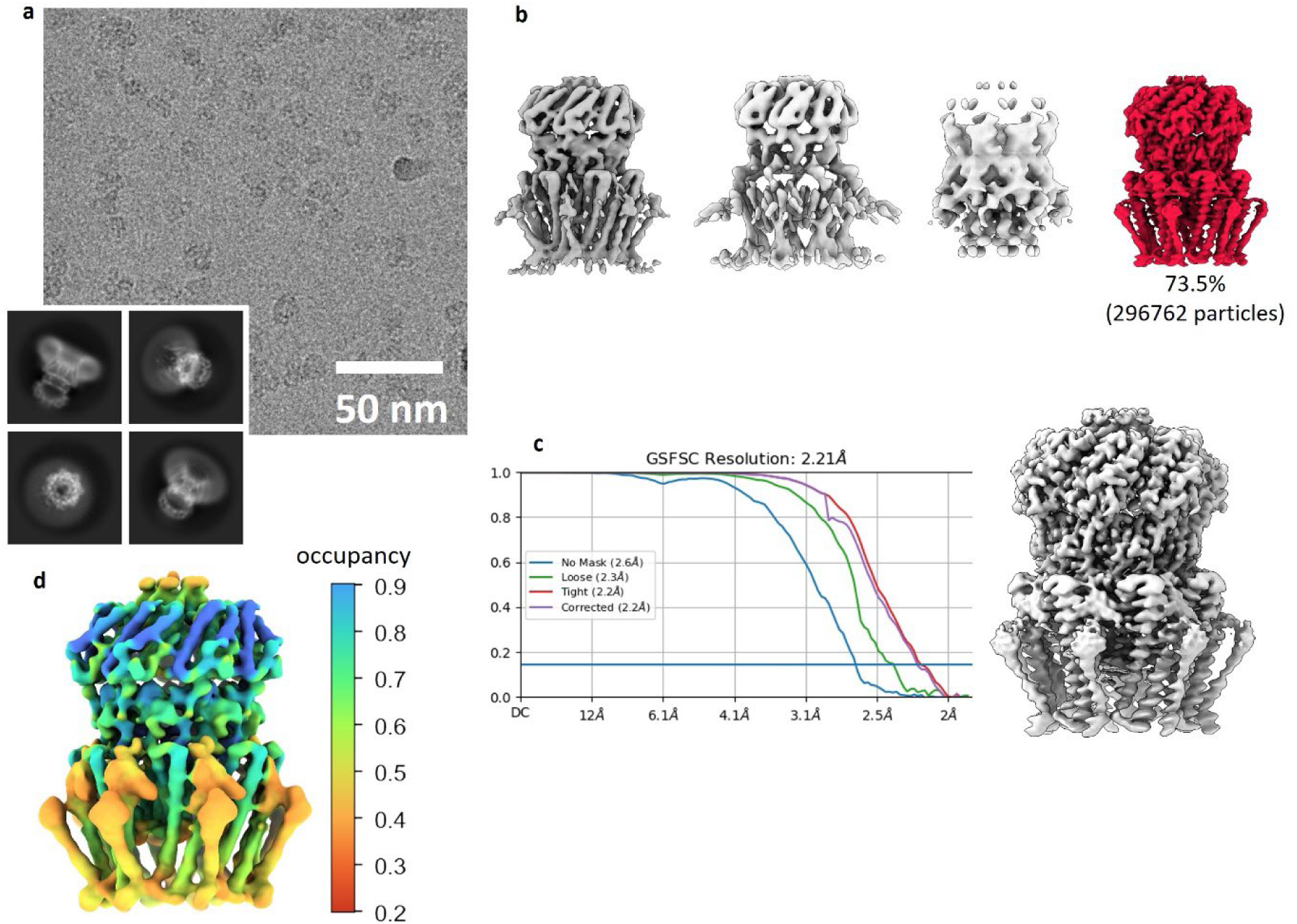
Cryo-EM analysis of closed WT-YnaI purified under condition I+. **a** One representative micrograph of the data collection is shown and four selected 2D class averages. The edge of the box corresponds to 24.2 nm. **b** A heterogenous refinement revealed one major class (red) that was further processed. **c** A non-uniform refinement resulted in a map at 2.2 Å resolution. **d** Occupancy estimation of the map was performed with OccuPy, showing lower values for the outer cytosolic loop and the helix TM(-2).

**Figure S4:**
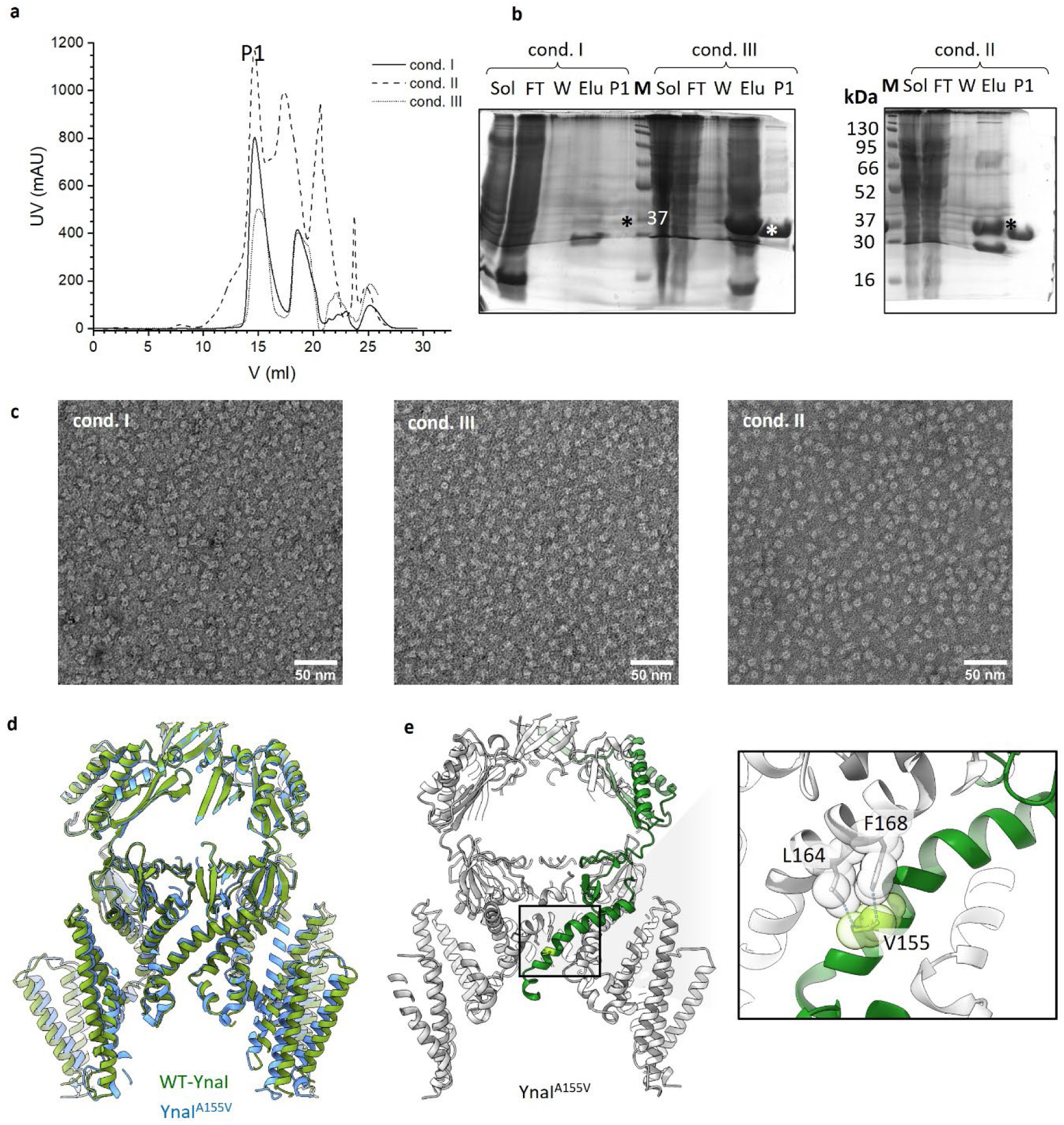
Purifications of YnaI^A155V^ under the different conditions. YnaI^A155V^ was purified under different conditions. **a** The size exclusion chromatograms are shown for YnaI purified under condition I+ (solid line), II (dashed line), and III (dotted line). The peak at 15 ml (P1) is the desired heptamer. **b** SDS-PAGES were done to monitor the purification. For all three purification approaches, samples were chosen from the supernatant after solubilisation (Sol), IMAC flowthrough (FL), IMAC wash (W), IMAC protein-containing elution fraction (Elu), and HPLC heptamer peak (P1), as well as a Marker (M). The signals from YnaI at the height of the 37 kDa marker signal are highlighted with an asterisk. **c** The P1 fraction was pooled and concentrated to the final concentration needed for subsequent cryo-EM preparation. Before, a dilution (1:2000-1:3000) of the concentrated sample was negatively stained and imaged for quality assessment. **d** The models of open wt-YnaI (green) and open YnaI^A155V^ (blue) are superposed. For clarity, only a central slice is shown. There are only minor differences in the vestibule, the pore, and the inner sensor paddle helices TM2 and TM1. However, the outer sensor paddle helices TM(-2) and TM(-1) exhibit slightly larger backbone variations. **e** A slice through YnaI^A155V^ is shown with one subunit depicted in green with the side chains of the substitutive Val155 coloured in light green. The black box is enlarged shown with the artificially introduced extra interaction between the Cγ atoms of Val155 with the side chains of Leu164 and Phe168. For better illustration, transparent van-der-Waals-radii are additionally shown for the mentioned residues.

**Figure S5:**
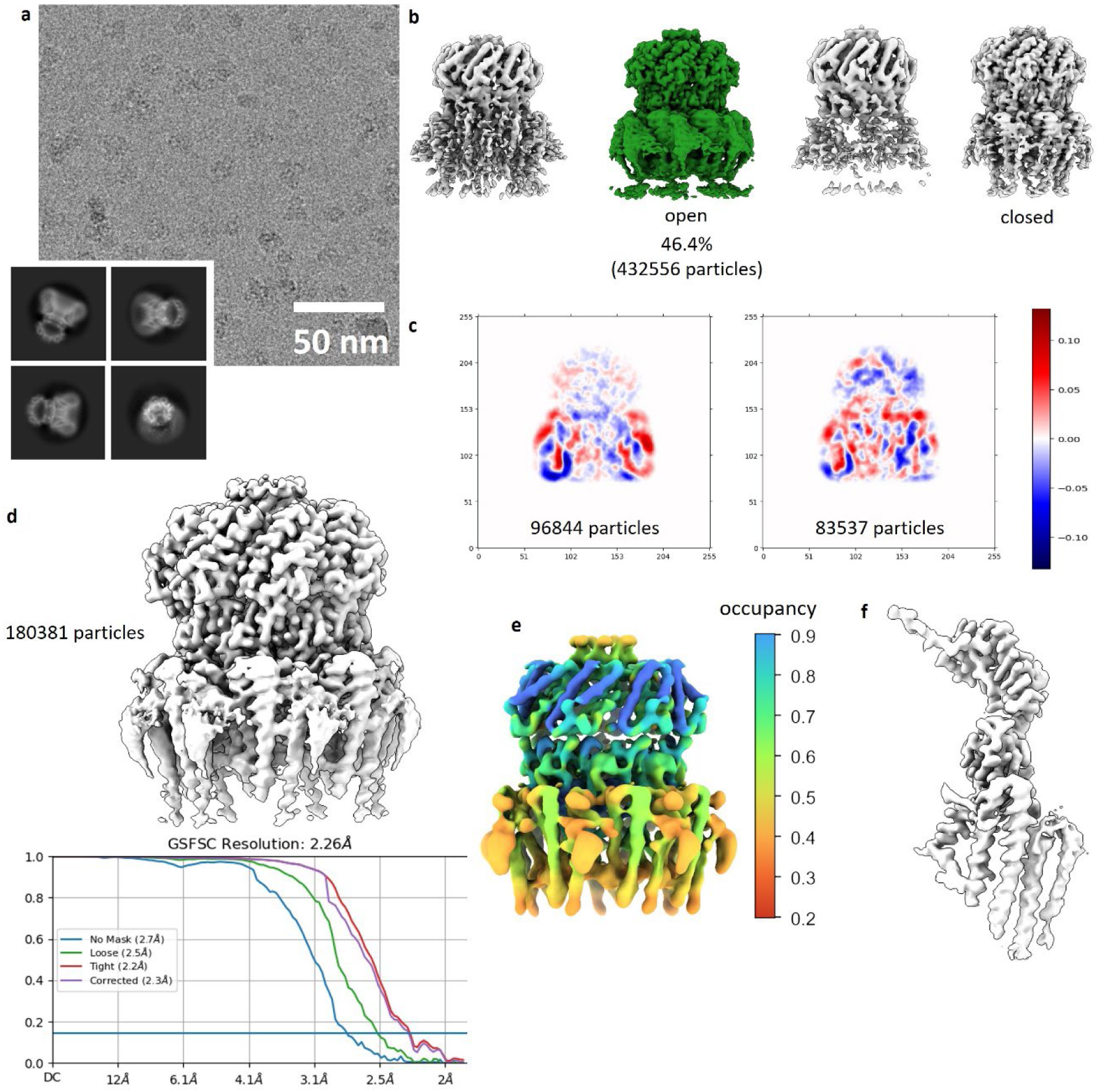
Cryo-EM analysis of open WT-YnaI. **a** One representative micrograph of the data collection is shown and four well represented 2D class averages. The edge of the box corresponds to 24.2 nm. **b** A heterogenous refinement revealed one open class (green) corresponding to approx. 46 % of all particles. This class was further processed. **c** A 3D variability analysis resulted in two clusters that performed best. **c** These were joined, and a non-uniform refinement resulted in a map at 2.3 Å resolution. **e** Occupancy estimation of the map was performed with OccuPy, showing least values for the outer cytosolic loop and the helix TM(-2). The scale bar on the right depicts the occupancy values. **f** To improve resolution and map quality of helix TM(-2), data was imported into Relion 5 and a refinement with C7 symmetry relaxation and blush denoising was conducted. The resulting density of the best resolved subunit is shown.

**Figure S6:**
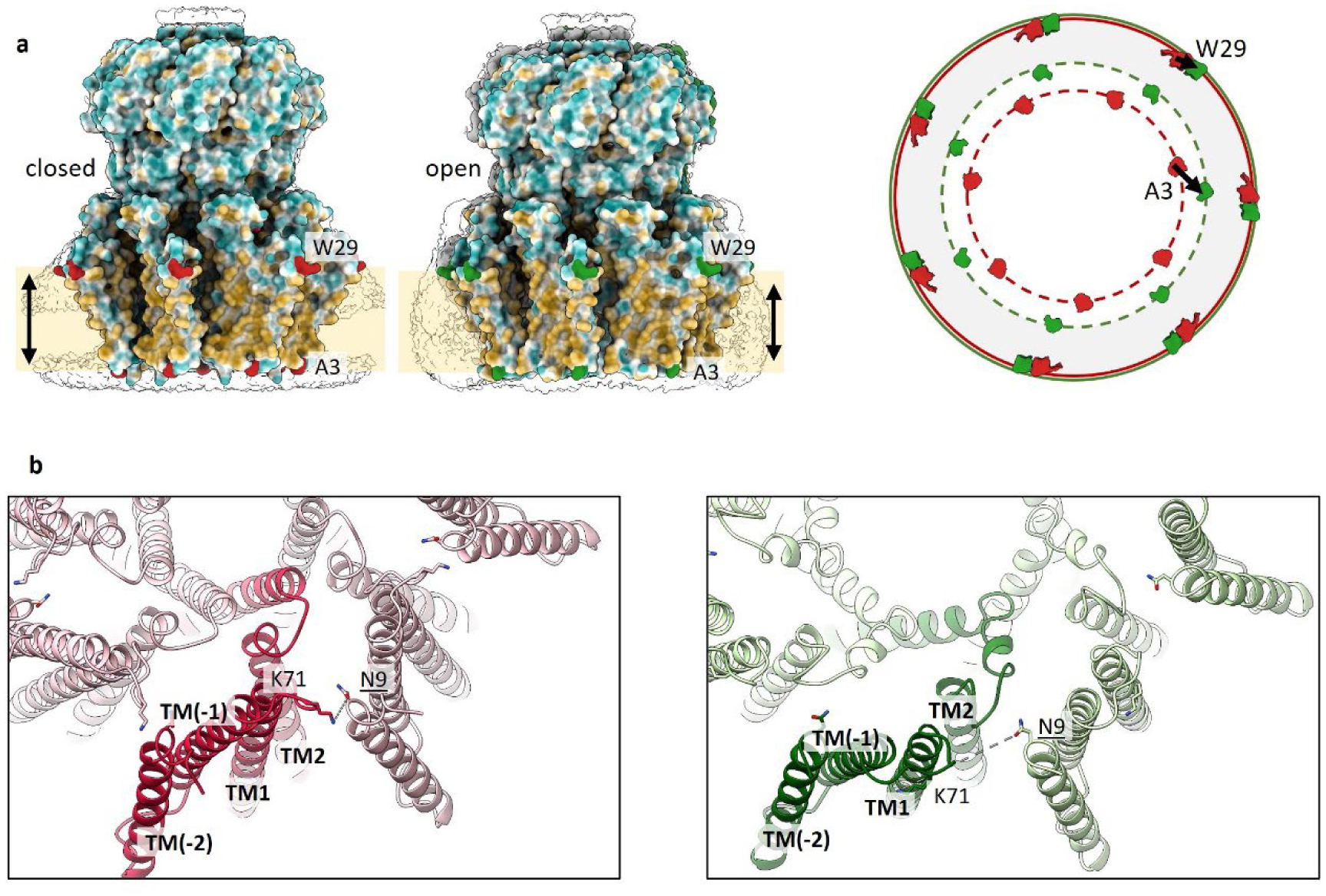
Comparison of the area change of the TM part of closed and open YnaI. **a** On the left, the surfaces of closed and open YnaI are coloured by their hydrophobicity. The significant residues Trp29, which marks the membrane interface at the cytosolic leaflet, and Ala3, which sits at the periplasmic side, are colored in red (closed YnaI) and green (open YnaI), respectively. In white, the map density is shown at a low threshold, highlighting the position of the detergent micelles. The yellow bars and black arrows depict the approximate membrane position and thickness. On the left, the highlighted residues are viewed from the periplasmic side, showing that the diameter of the channel at the cytosolic side (solid lines; red – closed; green – open) does not change upon opening, but at the periplasmic side (dotted lines). **b** YnaI is viewed from the periplasmic side. One subunit of closed (pink) and open (light green) YnaI is highlighted in a darker colour, and the TM helices are labelled. As only inter-subunit interaction in the paddles, Lys71 and Asn9 interact via H bonds in the closed state. In the open state, the paddles are radially relocated as rigid body. This results in further spacing of the paddles, and the H bond cannot be maintained.

**Figure S7:**
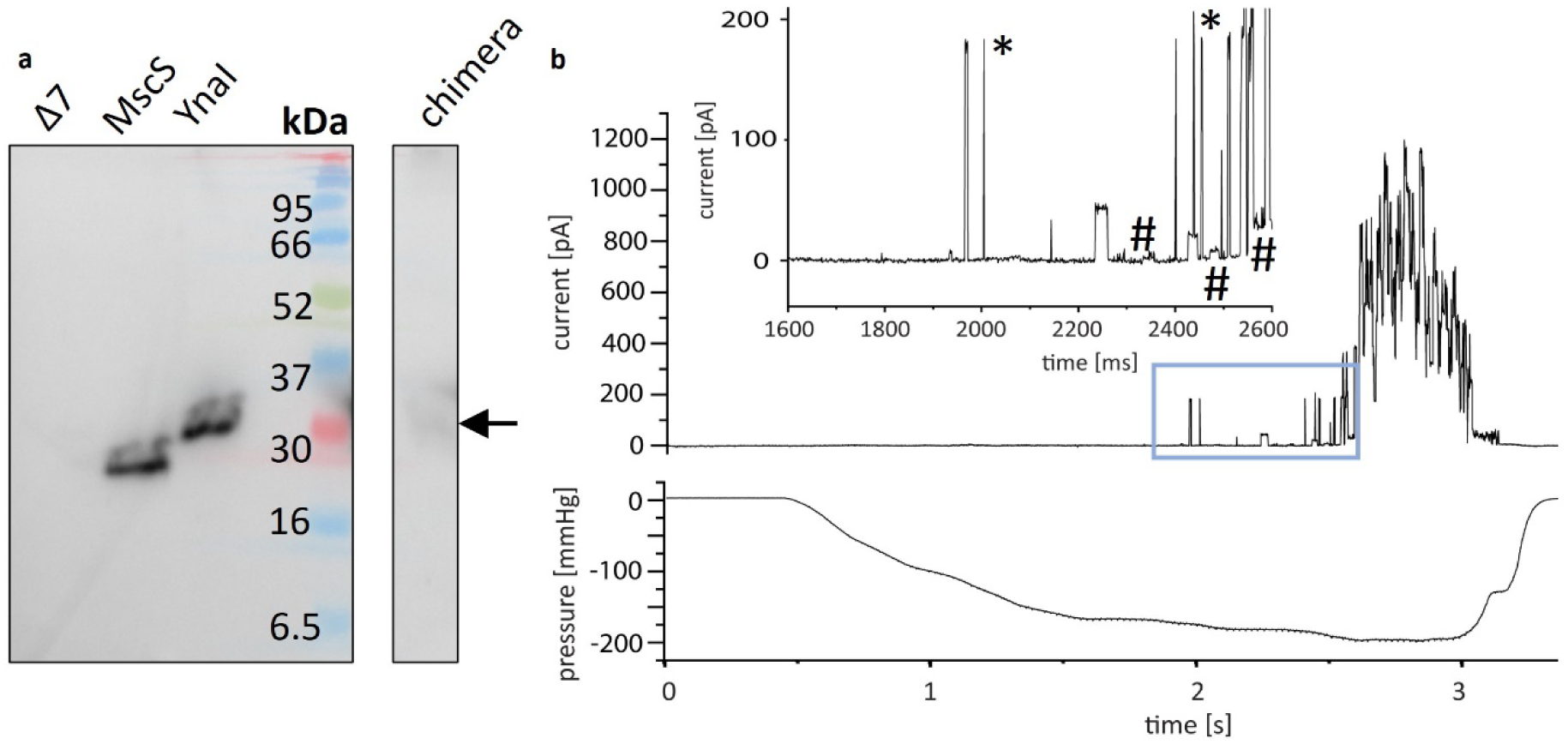
Functional analysis of a YnaI-MscS chimera. **a** The whole cell Western blot shows a weak overexpression of the chimera (black arrow) with a signal at a similar molecular weight as YnaI. **b** A current trace from membrane patches from *E. coli* MJF429-derived giant spheroplasts expressing the YnaI-MscS chimera is shown at the top and the corresponding pressure profile at the bottom. The membrane potential was +40 mV. The chimera opens at a similarly high pressure as MscL (*) and exhibits a conductance of ≥ 1 nS. Predominantly flickery or short openings are observable. The blue box is shown enlarged with the activities of the chimera labelled with #.

**Figure S8:**
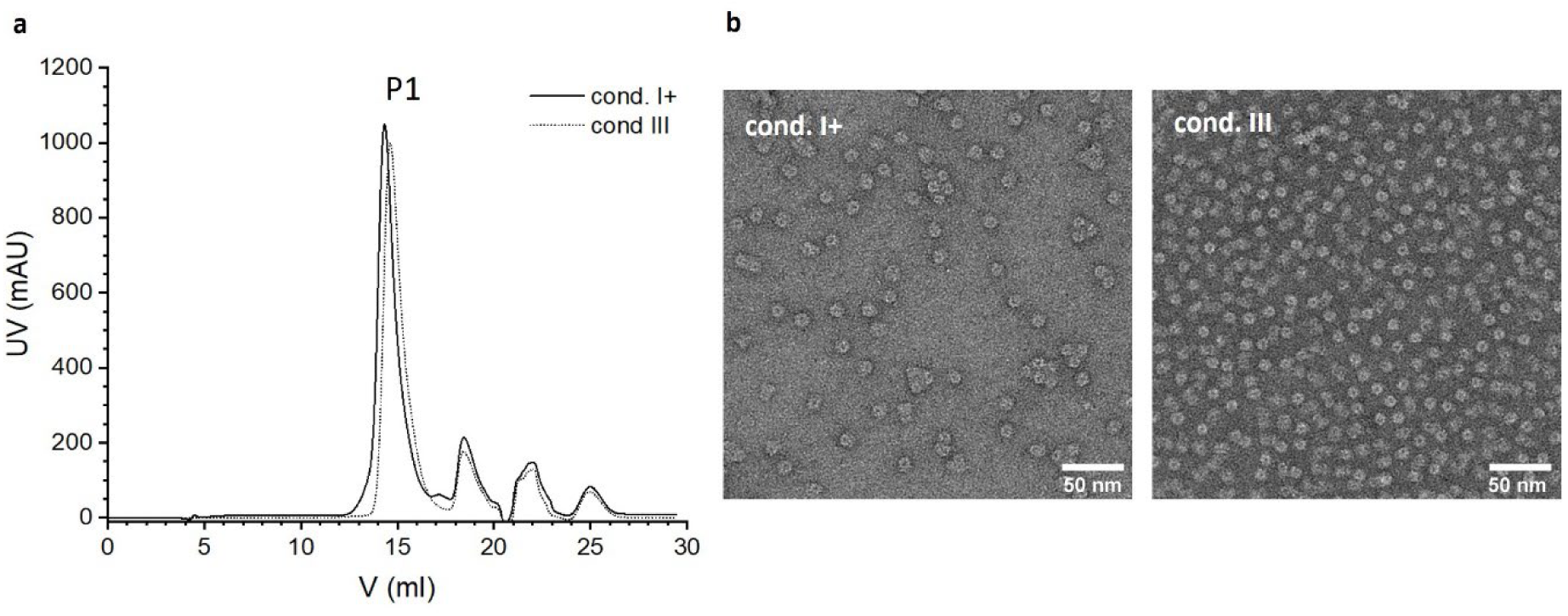
**Purification of closed and open YnaI-MscS chimera.** The YnaI-MscS chimera was purified under conditions I+ and III. **a** The size exclusion chromatograms are shown for YnaI purified under condition I+ (solid line) and III (dotted line) , and the peak at 15 ml (P1) is the desired heptamer. **b** The P1 fraction was pooled and concentrated to the final concentration needed for subsequent cryo-EM preparation. Before, a dilution (1:3000) of the concentrated sample was negatively stained and imaged for quality assessment.

**Figure S9:**
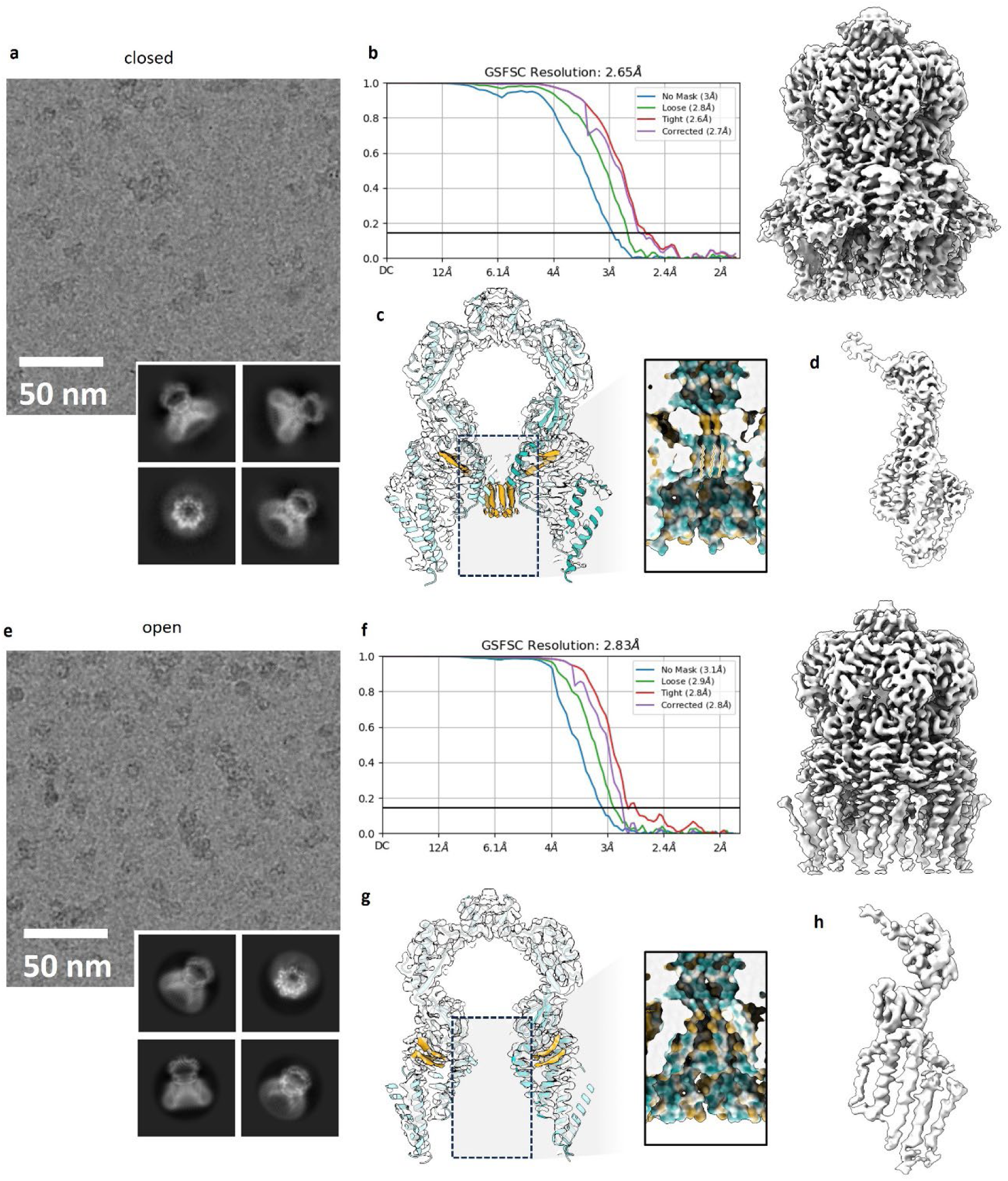
**Cryo-EM analysis of the closed and open YnaI-MscS chimera.** The cryo-EM analysis is shown for both the closed and open chimera. **a** A representative micrograph of the data collection of the chimera purified under condition I+ is shown and four well represented 2D class averages. The edge of the box corresponds to 24.2 nm. **b** The final map had a resolution of 2.7 Å resolution. **c** In a central slice, the model is coloured in light blue with one subunit darker, and lipid densities are depicted in orange. The insert in the box is shown enlarged with the model surface according to its hydrophobicity. Pore lipids are highlighted with a white silhouette. **d** To improve resolution and map quality of helix TM(-2), data was imported into Relion 5 and a refinement with C7 symmetry relaxation and blush denoising was conducted. The resulting density of the best resolved subunit is shown. **e** A representative micrograph of the data collection of the chimera purified under the condition III is shown and four well represented 2D class averages. The edge of the box corresponds to 24.2 nm. **f** The final map had a resolution of 2.8 Å resolution, but the transmembrane part is notably worse resolved compared to the closed chimera. **g** In a central slice, the model is coloured in light blue with one subunit darker, and lipid densities are depicted in orange. The insert in the box is shown enlarged with the model surface according to its hydrophobicity. **h** To improve resolution and map quality of helix TM(-2), data was imported into Relion 5 and a refinement with C7 symmetry relaxation and blush denoising was conducted. The resulting auto-sharpened density of the best resolved subunit is shown. The helical backbones of all helices are clearly visible, yet the density of the outermost two helices is not good enough to identify side chains.

**Table 1a:**
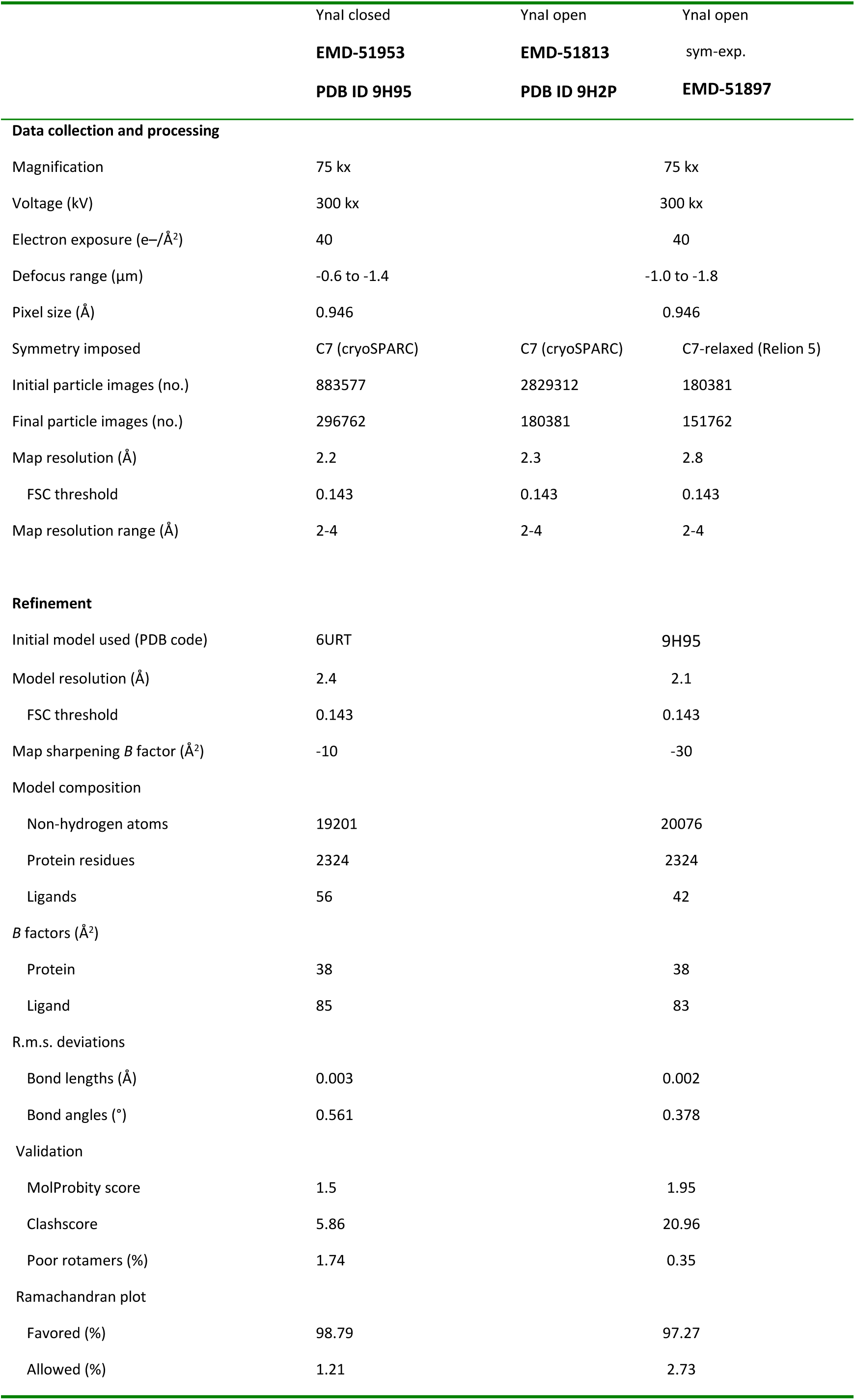
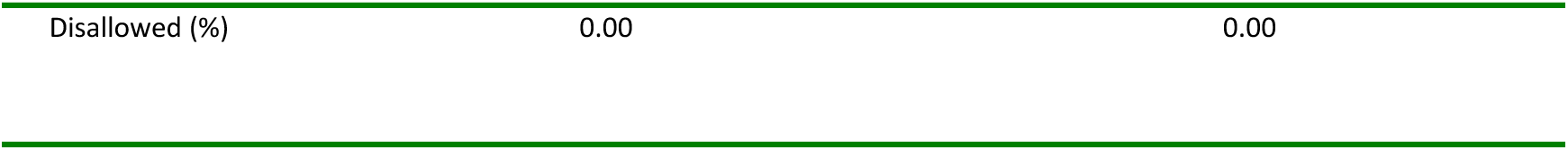
Cryo-EM data collection, refinement and validation statistics for closed and open WT-YnaI.

|  | Ynal closed | Ynal open | Ynal open |
| --- | --- | --- | --- |
|  | **EMD-51953** | **EMD-51813** | sym-exp. |
|  | **PDB ID 9H95** | **PDB ID 9H2P** | **EMD-51897** |
| **Data collection and processing** |  |  |  |
| Magnification | 75 kx |  | 75 kx |
| Voltage (kV) | 300 kx |  | 300 kx |
| Electron exposure (e-/Å<sup>2</sup>) | 40 |  | 40 |
| Defocus range (μm) | -0.6 to -1.4 |  | -1.0 to -1.8 |
| Pixel size (Å) | 0.946 |  | 0.946 |
| Symmetry imposed | C7 (cryoSPARC) | C7 (cryoSPARC) | C7-relaxed (Relion 5) |
| Initial particle images (no.) | 883577 | 2829312 | 180381 |
| Final particle images (no.) | 296762 | 180381 | 151762 |
| Map resolution (Å) | 2.2 | 2.3 | 2.8 |
| FSC threshold | 0.143 | 0.143 | 0.143 |
| Map resolution range (Å) | 2-4 | 2-4 | 2-4 |
| **Refinement** |  |  |  |
| Initial model used (PDB code) | 6URT |  | 9H95 |
| Model resolution (Å) | 2.4 |  | 2.1 |
| FSC threshold | 0.143 |  | 0.143 |
| Map sharpening *B* factor (Å<sup>2</sup>) | -10 |  | -30 |
| Model composition |  |  |  |
| Non-hydrogen atoms | 19201 |  | 20076 |
| Protein residues | 2324 |  | 2324 |
| Ligands | 56 |  | 42 |
| *B* factors (Å<sup>2</sup>) |  |  |  |
| Protein | 38 |  | 38 |
| Ligand | 85 |  | 83 |
| R.m.s. deviations |  |  |  |
| Bond lengths (Å) | 0.003 |  | 0.002 |
| Bond angles (°) | 0.561 |  | 0.378 |
| Validation |  |  |  |
| MolProbity score | 1.5 |  | 1.95 |
| Clashscore | 5.86 |  | 20.96 |
| Poor rotamers (%) | 1.74 |  | 0.35 |
| Ramachandran plot |  |  |  |
| Favored (%) | 98.79 |  | 97.27 |
| Allowed (%) | 1.21 |  | 2.73 |

|  |  |  |
| --- | --- | --- |
| Disallowed (%) | 0.00 | 0.00 |

**Table 1b:**
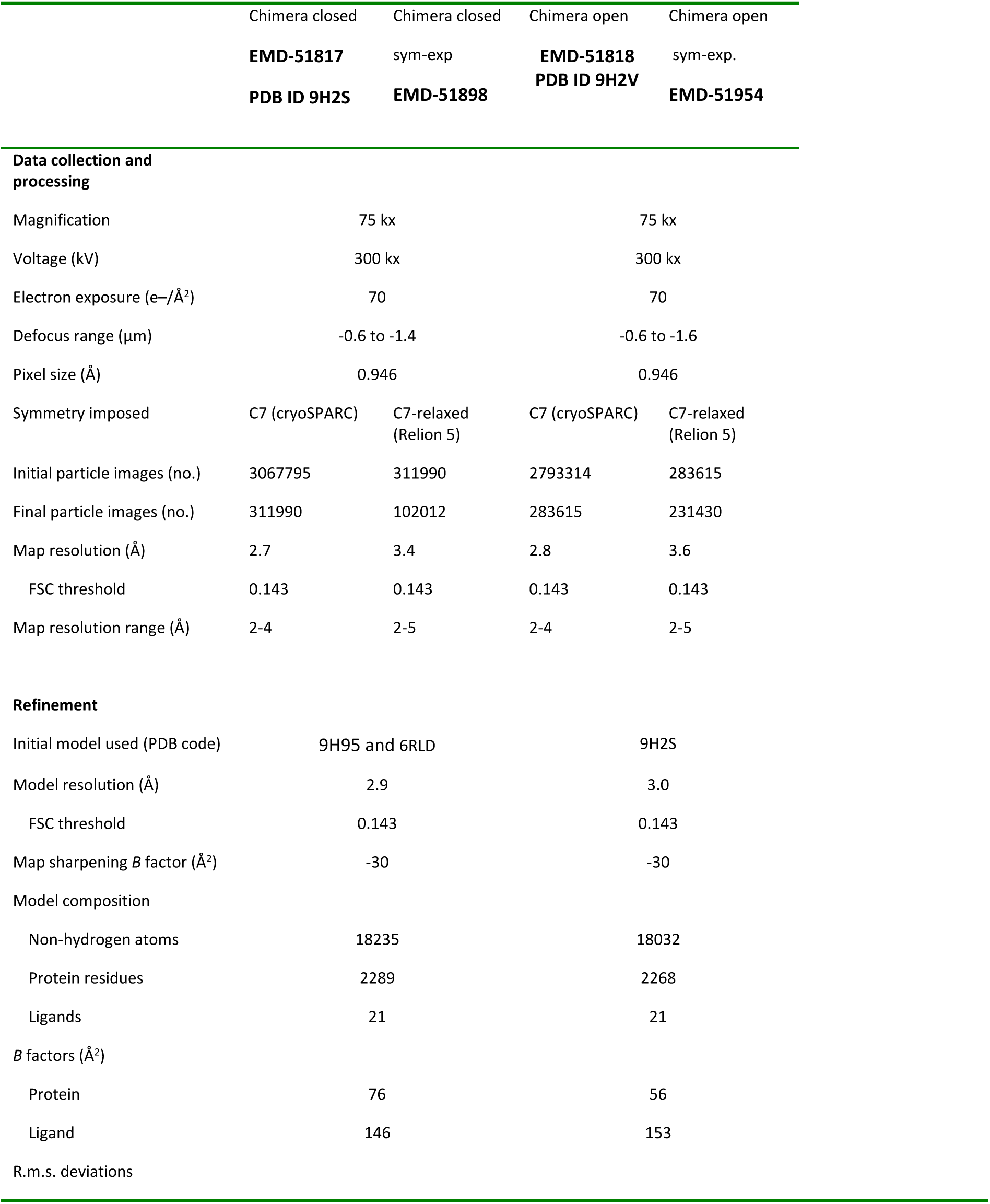
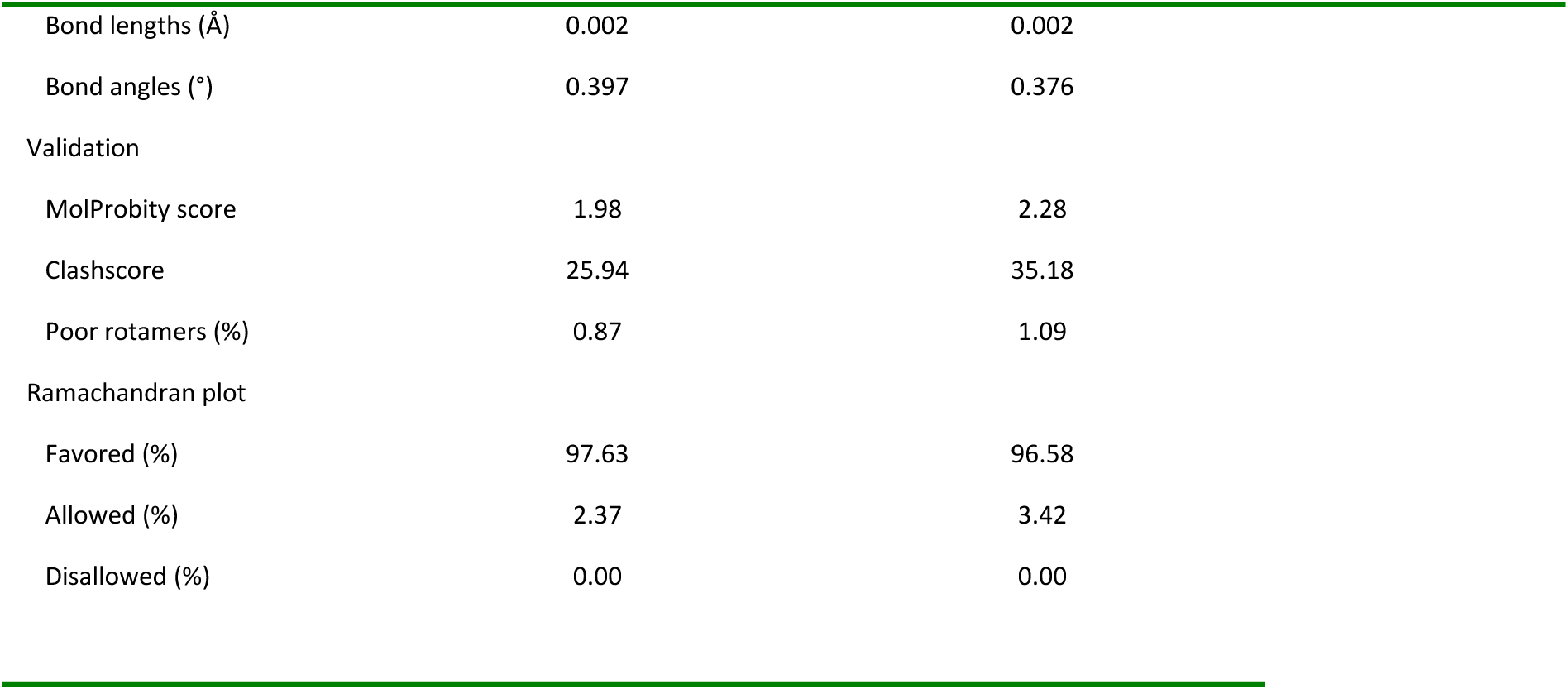
Cryo-EM data collection, refinement and validation statistics for closed and open YnaI-MscS chimera.

|  | Chimera closed | Chimera closed | Chimera open | Chimera open |
| --- | --- | --- | --- | --- |
|  | EMD-51817 | sym-exp | EMD-51818 | sym-exp. |
|  | PDB ID 9H2S | EMD-51898 | PDB ID 9H2V | EMD-51954 |
| Data collection and processing |  |  |  |  |
| Magnification | 75 kx |  | 75 kx |  |
| Voltage (kV) | 300 kx |  | 300 kx |  |
| Electron exposure (e-/Å²) | 70 |  | 70 |  |
| Defocus range (µm) | -0.6 to -1.4 |  | -0.6 to -1.6 |  |
| Pixel size (Å) | 0.946 |  | 0.946 |  |
| Symmetry imposed | C7 (cryoSPARC) | C7-relaxed (Relion 5) | C7 (cryoSPARC) | C7-relaxed (Relion 5) |
| Initial particle images (no.) | 3067795 | 311990 | 2793314 | 283615 |
| Final particle images (no.) | 311990 | 102012 | 283615 | 231430 |
| Map resolution (Å) | 2.7 | 3.4 | 2.8 | 3.6 |
| FSC threshold | 0.143 | 0.143 | 0.143 | 0.143 |
| Map resolution range (Å) | 2-4 | 2-5 | 2-4 | 2-5 |
| Refinement |  |  |  |  |
| Initial model used (PDB code) | 9H95 and 6RLD |  | 9H2S |  |
| Model resolution (Å) | 2.9 |  | 3.0 |  |
| FSC threshold | 0.143 |  | 0.143 |  |
| Map sharpening B factor (Å²) | -30 |  | -30 |  |
| Model composition |  |  |  |  |
| Non-hydrogen atoms | 18235 |  | 18032 |  |
| Protein residues | 2289 |  | 2268 |  |
| Ligands | 21 |  | 21 |  |
| B factors (Å²) |  |  |  |  |
| Protein | 76 |  | 56 |  |
| Ligand | 146 |  | 153 |  |
| R.m.s. deviations |  |  |  |  |

|  |  |  |
| --- | --- | --- |
| Bond lengths (Å) | 0.002 | 0.002 |
| Bond angles (°) | 0.397 | 0.376 |
| Validation |  |  |
| MolProbity score | 1.98 | 2.28 |
| Clashscore | 25.94 | 35.18 |
| Poor rotamers (%) | 0.87 | 1.09 |
| Ramachandran plot |  |  |
| Favored (%) | 97.63 | 96.58 |
| Allowed (%) | 2.37 | 3.42 |
| Disallowed (%) | 0.00 | 0.00 |

